# Extensive richness and novel taxa of sulfoquinovose-degrading bacteria in the cow rumen

**DOI:** 10.1101/2025.05.20.655074

**Authors:** Julia Krasenbrink, Song-Can Chen, Tomohisa Sebastian Tanabe, Hüseyin Sarikeçe, Pleun Meurs, Sabrina Borusak, Rahul Samrat, Jay Osvatic, Joana Séneca, Bela Hausmann, Daan R. Speth, Evelyne Selberherr, Wolfgang Wanek, David Schleheck, Marc Mussmann, Alexander Loy

**Affiliations:** University of Vienna, Centre for Microbiology and Environmental Systems Science, Division of Microbial Ecology, Vienna, Austria; University of Vienna, Doctoral School in Microbiology and Environmental Science, Vienna, Austria; State Key Laboratory of Soil Pollution Control and Safety, Zhejiang University, Hangzhou, China; MOE Key Laboratory of Environment Remediation and Ecological Health, College of Environmental and Resource Sciences, Zhejiang University, Hangzhou, China; Department of Biology, University of Konstanz, Konstanz, Germany; Konstanz Research School Chemical Biology, University of Konstanz, Konstanz, Germany; University of Vienna, Centre for Microbiology and Environmental Systems Science, Division of Terrestrial Ecosystem Research, Vienna, Austria; Joint Microbiome Facility of the Medical University of Vienna and the University of Vienna, Vienna, Austria; Division of Clinical Microbiology, Department of Laboratory Medicine, Medical University of Vienna, Vienna, Austria; Unit of Food Microbiology, Institute of Food Safety, Food Technology and Veterinary Public Health, University of Veterinary Medicine, Vienna, Austria

**Author notes:** Corresponding author: Alexander Loy. These authors contributed equally: Song-Can Chen and Tomohisa Sebastian Tanabe.

**Keywords:** Sulfoquinovose, sulfur cycle, organosulfur, sulfolipid, gut microbiome, rumen, marine sediment, functional gene, sulfoquinovosidase

## Abstract

Sulfoquinovose (SQ), a sulfonated sugar derived from the thylakoid membrane lipid sulfoquinovosyl diacylglycerol (SQDG), is abundant in photosynthetic organisms and plays a key role in global sulfur cycling. Its degradation in nature is mediated by specialized bacteria, many of which rely on the enzyme sulfoquinovosidase (YihQ) to release SQ from SQDG. Despite its ecological importance, the diversity and functional roles of SQ-degrading microorganisms remain poorly characterized in natural environments. Here, we developed a *yihQ*-targeted amplicon sequencing approach to investigate the richness and distribution of SQ-degrading bacteria across selected environments, including marine sediments and the mammalian gut. We revealed particularly high richness of *yihQ*-containing microorganisms in cow rumen, far exceeding that observed in human and mouse gut microbiomes, suggesting an important role of SQ metabolism in ruminant digestion. Anaerobic microcosm experiments with SQ-amended rumen fluid revealed cooperative microbial degradation of SQ to sulfide via isethionate cross-feeding. Amplicon sequencing and genome-resolved metagenomics identified novel uncultured SQ-degrading taxa, including members of *Caproiciproducens* (*Acutalibacteraceae*), *Limivicinus* (*Oscillospiraceae*), and *Sphaerochaetaceae*, which encode the sulfo-transketolase pathway, along with *Mailhella* (*Desulfovibrionaceae*), a likely isethionate-respiring bacterium. This study presents the first functional gene-based assay for tracking environmental *yihQ* diversity, highlights SQ degradation as a central metabolic process in the cow rumen, describes novel SQ-metabolizing bacteria, and advances understanding of sulfur physiology in complex microbial communities.

## Introduction

Sulfoquinovose (SQ; 6-deoxy-6-sulfo-D-glucose) is a sulfonated hexose, which is ubiquitous in the environment. Approximately 10 billion tons of SQ are produced globally per year, comparable to the amino acids methionine and cysteine. SQ is an integral component of the biogeochemical sulfur cycle and the human and animal diet [1,2]. While free SQ appears to be rare in nature, it is primarily present as the head group of the sulfolipid sulfoquinovosyl diacylglycerol (SQDG) in the biomass of photosynthetic organisms such as plants, algae, and cyanobacteria [2–6]. SQDG resides in the thylakoid membrane, where it supports the activity and structural integrity of photosystem II [2,7,8].

To date, several pathways for SQ degradation have been identified in bacteria. Some pathways require inter-species transfer of organosulfonates for complete SQ mineralization to sulfate or hydrogen sulfide [9–15]. In contrast, the SQ monooxygenase pathway in *Agrobacterium tumefaciens* and the *Roseobacter* clade and the SQ dioxygenase pathway in *Marinomonas ushuaiensis* aerobically cleave the C-S bond of SQ and directly catabolize SQ to sulfite and a deoxy-hexose [16–18]. Since SQDG is the main natural source of SQ, all known pathways require the enzymatic release of SQ from SQDG or sulfoquinovosyl glycerol [4,19]. In five of the six physiologically and biochemically characterized SQ degradation pathways, this cleavage is catalyzed by sulfoquinovosidases of the glycoside hydrolase family 31 [4,10,11,13,14,16,17,19–21]. These include YihQ in *Escherichia coli* [10], *Akalicoccus urumqiensis* (formerly *Bacillus urumqiensis*) [14], *Clostridium* sp. [14], and *Novosphingobium aromaticivoran*s [14], PpSQ1_00094 in *Pseudomonas putida* [11], SftG in *Bacillus aryabhattai* [13], SqvC in *Priestia megaterium* (formerly *Bacillus megaterium*) [21], and SmoI in *A. tumefaciens* [16]. These sulfoquinovosidases have conserved structural and functional features and form a monophyletic orthologous group in the glycoside hydrolase family 31 tree [4,19]. We therefore refer to members of this orthologous group as YihQ sulfoquinovosidases. YihQ from *E. coli* and *A. tumefaciens* hydrolyzed both SQDG and sulfoquinovosyl glycerol *in vitro* [4,19]. *E. coli* cultures supplemented with sulfoquinovosyl glycerol or glucose grew to the same optical density. In these growth experiments sulfoquinovosyl glycerol was metabolized faster than glycerol and free sulfoquinovose, but later than glucose. Growth rates of cultures with sulfoquinovosyl glycerol were also higher than those of cultures grown solely with SQ or glycerol alone, indicating that sulfoquinovosyl glycerol is the preferred substrate for *E. coli* [19].

Genes of YihQ are widespread among bacteria that encode SQ-degradation pathways. Some SQ-degrading bacteria, such as *Roseobacter* spp., *Arthrobacter* spp., and *Marinomonas* sp., lack a *yihQ* homolog [17,18,22]. However, a new family of NAD^+^-dependent sulfoquinovosidases was recently discovered and is widely distributed in bacteria that were previously thought to lack a sulfoquinovosidase [23].

SQDG and SQ are integral parts of the sulfur cycle, yet the diversity of SQ-degrading microorganisms across different environments remains largely unexplored, with studies to date limited to human and mouse gut microbiomes and seawater [18,24,25]. The presence of *yihQ* homologs in five of the six SQ degradation pathways suggests that this gene represents a suitable functional marker for many SQDG/SQ-degrading microorganisms. To this end, we developed two *yihQ*-targeted degenerate primer sets and evaluated their use for analyzing the *yihQ* sequence diversity in samples from various environments (soil, marine sediments) and host-associated gut microbiomes (human, cow, and mouse). Furthermore, the *yihQ* amplicon sequencing approach was combined with 16S rRNA gene amplicon sequencing, metagenomics, and metabolite analysis to study, for the first time, SQ degradation and the involved microorganisms and pathways in cow rumen fluid microcosms. We show that the SQ-degrading community in the cow rumen differs from those in the mouse and human gut by comprising several hundred distinct *yihQ* sequences, consisting of undescribed species, and predominantly utilising the sulfo-transketolase pathway for SQ degradation to isethionate.

## Materials and Methods

Supplementary Materials and Methods provide further details on the methods described below.

### Design of *yihQ*-targeted PCR assays

A YihQ reference database was created using sequences of biochemically validated YihQ sulfoquinovosidases and homologs of functionally different enzymes of the glycoside hydrolase family 31 as queries for Blast against the KEGG prokaryotes database [26] (Table S1). An alignment of representative sequences was used for calculation of a maximum-likelihood YihQ tree (Fig. S1) and for the design of degenerate *yihQ*-targeted primers. Genomic DNA of the *yihQ*-encoding strains *Escherichia coli* K12, *Agathobacter rectalis* A1-86, and *Enterocloster clostridioformis* YL32 were used as positive controls for PCR. DNA of *Segatella copri* DSM 18205 and ultrapure water served as non-target control and negative control, respectively. Primer pairs targeting different *yihQ* regions were initially tested using pure culture DNA for temperature gradient PCR and further optimized by touchdown PCR. The optimized PCR program (Table S2) was subsequently tested with DNA from several environmental and intestinal samples.

### Sampling and ethics

Marine coastal surface sediment samples were collected from Canada (Vancouver: 49°17’57.7"N, 123°10’34.6"W), Italy (Fetovia: 42°43’57.2"N, 10°9’15.6"E), Mauritania (Dakhlet Nouadhibou: 20°36’52.7"N, 16°30’35.9"W), and Australia (Casuarina Beach: 12°21’39.96"S, 130°51’56.7"E). A soil sample was collected from the acidic peatland Schlöppnerbrunnen II, Germany (50°07ʹ54.8ʺN, 11°52ʹ51.8ʺE), as previously described [27]. A human fecal sample mix was created by pooling freshly collected feces from five adult volunteers. Sampling and microbiota analyses of human feces was approved by the ethics commission of the University of Vienna (reference numbers 00161 and 00714). A mouse gut content sample mix was created by pooling small intestinal and cecal contents from eight conventional C57BL/6 wildtype mice (4 females, 4 males, aged 8–10 weeks). Mice were housed under specific-pathogen-free conditions in compliance with the Federation of European Laboratory Animal Science Associations guidelines, with *ad libitum* access to a standard chow diet (Ssniff V1534, Ssniff, Germany) and water. Rumen fluid was collected at an International Featured Standard-Food certified slaughterhouse in Lower Austria. Cows originated from the same farm, were fed hay and grass silage *ad libitum*, with a maximum of 0.5 kg/day of commercial concentrate (KuhKorn PLUS Energie; Garant-Tiernahrung GmbH, Austria), and had free access to water, which is common practice in the region. Post-mortem, the mechanical isolation procedure and gastrointestinal tract removal were performed by licensed veterinarians following standard operating protocols to minimize contamination and preserve microbial composition. A volume of 50 ml of rumen fluid was obtained from each rumen under controlled hygienic conditions. Samples were immediately transported to the laboratory and processed. During the first collection, a sample was obtained from one cow and used for quantifying SQDG and for *yihQ*-amplicon sequencing alongside with other environmental samples. During the second collection, one sample each was obtained from three cows and was used for the rumen fluid microcosm experiment. As the mouse gut and rumen fluid samples were obtained post-mortem, no ethical approval was required for this study.

### Rumen fluid microcosms

Anoxic rumen fluid microcosms supplemented with 10 mM SQ (MCAT GmbH, Germany) were used to investigate SQ metabolism by the cow gut microbiome (Supplementary Materials & Methods). Microcosms were subsampled over 168 hours for metabolite quantification and microbial community analyses.

### Metabolite analysis

Ultra-high-performance reverse phase liquid chromatography combined with mass spectrometry was used to quantify and identify all molecular species of SQDG in rumen fluid (Supplementary Materials & Methods). Sulfide was quantified colorimetrically as previously described [28]. Formate, succinate, acetate, propionate, butyrate, valerate, and lactate were quantified by capillary electrophoresis. SQ and its metabolites isethionate, 2,3-dihydroxypropane-1-sulfonate (DHPS), 3-hydroxypropane-1-sulfonate, 3-sulfolactate, and 3-sulfopropionate were quantified by liquid chromatography–mass spectrometry as described previously (Supplementary Materials & Methods) [29].

### 16S rRNA gene and *yihQ* amplicon sequencing and analyses

Genomic DNA from environmental, intestinal, and rumen fluid microcosm samples was isolated using the DNeasy PowerSoil Pro Kit (Qiagen, Austria). Amplicon sequencing on the MiSeq platform (V3, 600 cycles, Illumina, USA) was conducted by the Joint Microbiome Facility (University of Vienna, Medical University of Vienna) under project IDs JMF-2310-13 and JMF-2303-02. DNA from each environmental sample was sequenced in triplicates within the same sequencing run. A two-step PCR method [30,31] and primer pairs YIHQa (YIHQ1201Fa/YIHQ1526Ra) and YIHQb (YIHQ1201Fb/YIHQ1526Rb) were used to amplify and barcode an approximately 340 bp region of the *yihQ* gene (Fig. 1, Table S2). Samples from the rumen fluid microcosm experiment were analyzed by amplicon sequencing with primer pair YIHQb and with 16S rRNA gene-targeted primers for bacteria and archaea [30,31]. Amplicon sequence variants (ASVs) were inferred using DADA2 [32]. FASTQ reads 1 and 2 were trimmed at 220 nt and 150 nt, respectively, with allowed expected errors of 2. 16S rRNA gene-ASVs were classified using DADA2 and the SILVA 16S rRNA sequence database (release 138.1) [33]. A workflow was established for analysis of sequences obtained with *yihQ*-targeted primers, including removal of non-*yihQ* sequences and phylogenetic placement in the YihQ reference tree (Fig. S2, Supplementary Materials & Methods). Verified *yihQ* ASV sequences were further clustered into operational taxonomic units (OTUs) using vsearch [34] with a 90% sequence identity threshold. Alpha diversity metrics (observed number of ASVs and OTUs, Shannon diversity index, and Simpson diversity index) were calculated for subsampled (100, 1000, and 10000 reads) *yihQ* sequence libraries. Verified *yihQ* ASV sequences generated with the YIHQb amplicon sequencing assay were additionally classified using the GlobDB genome database (version r220) [35].

**Fig. 1.**
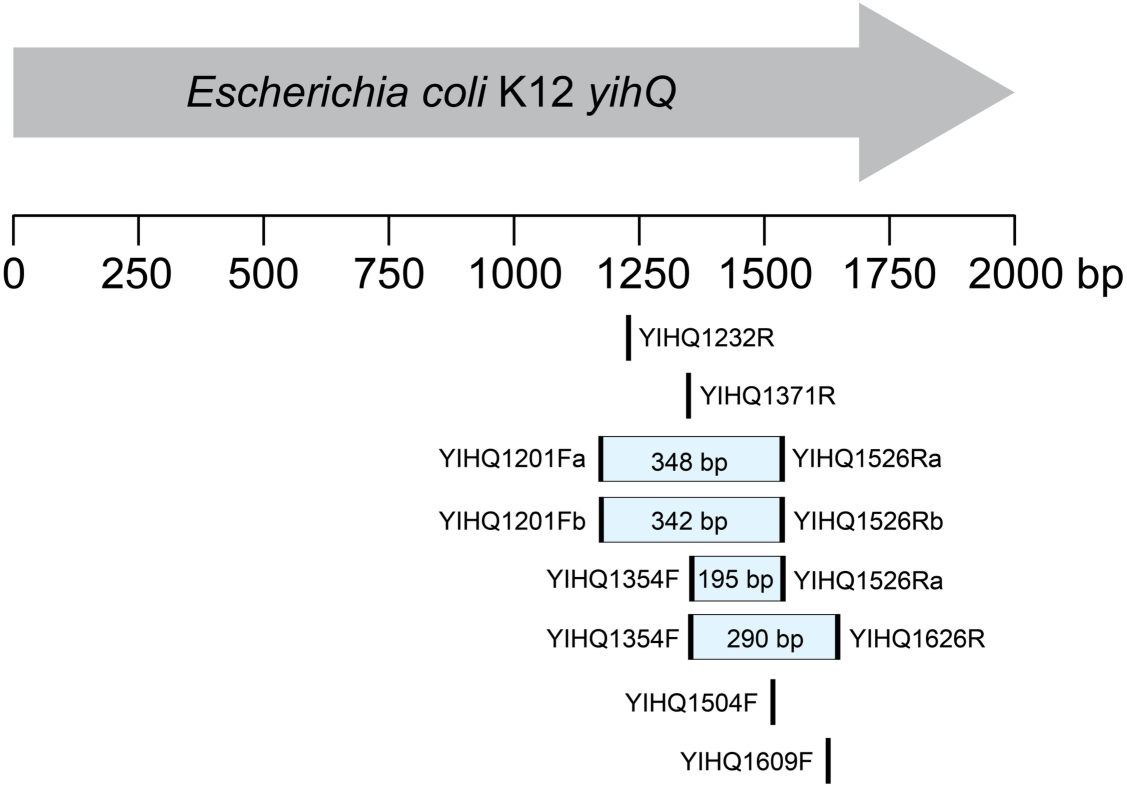
*yihQ*-targeted primers designed in this study. Primer names refer to the target gene *yihQ* and the binding position in the *yihQ* sequence of *E. coli* K12 (2037 bp, NCBI gene ID: 948376). Bars indicate the predicted length of the *yihQ* PCR product in *E. coli* K12. Primers without bars were not tested. bp = base pairs.

Significantly enriched 16S rRNA gene- and *yihQ-*OTUs/ASVs in SQ-supplemented rumen fluid microcosms were identified by DESeq2 (v1.38.3) as described previously [36]. Enriched ASVs/OTUs were taxonomically classified by Blast against the NCBI 16S rRNA sequence database and the GlobDB database [35,37].

### Metagenomics and genome analysis

Metagenomic sequencing was performed on a NovaSeq6000 (1/2 SP flow cell, 2×100 bp reads; Illumina, USA) with DNA extracts from SQ-amended (n = 2) and unamended control (n = 1) microcosms of rumen fluid, taken after 168 hours of incubation. Metagenome-assembled genomes (MAGs) were dereplicated and taxonomically classified using GTDB-Tk [38]. Average amino acid identity (AAI) was calculated using the Enveomics Collection [39]. Whole-genome average nucleotide identity (ANI) was calculated using FastANI (version 1.33) [40].

New Hidden Markov Models (HMMs) were generated for the detection of enzymes and transport proteins of sulfoquinovose, isethionate, and DHPS degradation pathways (Table S3). HMMs were integrated into HMSS2 [41], which was then used to identify sulfur metabolism genes in the recovered MAGs and GlobDB genomes (Supplementary Materials & Methods).

### Data and code availability

All sequence data generated in this project are available at NCBI under BioProject ID PRJNA1242727. The YihQ reference database, the amino acid alignment of YihQ reference sequences, the YihQ reference tree, and further information are available in Additional Files 1-7. HMMs for microbial organosulfur metabolism proteins are available at github.com/TSTanabe/HMSS2/tree/main/Hidden_Markov_Models.

## Results and Discussion

### Development of PCR assays for broad coverage of *yihQ* sequence diversity

Based on a curated YihQ sequence database and a YihQ reference tree (Fig. S1), we designed ten degenerate primers that target different conserved regions in the *yihQ* alignment (Fig. 1, Table S4). Of the four tested primer combinations, only two primer pairs, YIHQa (YIHQ1201Fa/YIHQ1526Ra) and YIHQb (YIHQ1201Fb/YIHQ1526Rb) resulted in PCR products of the expected size with DNA templates of three *yihQ*-containing strains in the temperature gradient PCR tests. Primer pairs YIHQa and YIHQb were further tested in a touchdown PCR approach for enhanced specificity. The final protocol for both primer pairs used an initial annealing temperature of 63°C, which decreased by 1°C per cycle over 10 cycles, followed by 20 cycles at 53°C. Subsequent magnesium chloride concentration tests showed that 4 mM magnesium chloride in the PCR buffer consistently produced the strongest bands for the target sequences, while the negative and non-target-DNA controls did not yield a visible PCR product (Table S2). Sanger sequencing of the PCR products confirmed specific amplification of the *yihQ* target gene in all three organisms.

The primers YIHQ1201Fb and YIHQ1526Rb were designed as shorter and less degenerate alternatives to YIHQ1201Fa and YIHQ1526Ra. Thus, primer pairs YIHQa and YIHQb amplify the same, approximately 340 bp region of *yihQ*, yet differ in degeneracy, predicted melting temperatures, and sequence coverage (Table S4), which may influence PCR amplification efficiency and specificity [42]. The YIHQa and YIHQb primer pairs perfectly match 81.8% and 89.8% of the *yihQ* sequences in the reference database, respectively. We thus applied and compared both primer pairs for amplification of DNA from various intestinal and environmental samples. These included pooled mouse gut content and human feces, which are known to harbor *yihQ*-containing bacteria [24,25], and rumen fluid, soil, and diverse marine sediments. The latter are environments that have not yet been investigated for their SQ degradation capacity. Both PCR assays successfully amplified a single band of the expected size with DNA from all intestinal samples. PCRs with peat soil DNA were negative. PCRs with primer pair YIHQa produced faint PCR products with DNA from Canadian marine sediment, but no PCR products with DNA from Italian, Mauritanian, and Australian marine sediments. In contrast, PCRs with primer pair YIHQb generated bands for all marine sediment samples.

### Optimized *yihQ* amplicon sequencing reveals highest *yihQ* richness in cow rumen

We optimized the YIHQa and YIHQb PCR assays for amplicon sequencing on the Illumina MiSeq platform. Both primer pairs contained a generic 16 bp 5’-head sequence for two-step PCR barcoding [30,31] (Table S2). The optimized PCR protocol was applied for amplification of DNA from the intestinal and environmental samples. Amplicons of the correct size were obtained with DNA from rumen fluid, mouse gut content, human feces, and Canadian sediment with both PCR assays. Additional amplicons with DNA from Mauritanian and Italian sediment were only obtained with the YIHQb assay, which suggests it has higher sensitivity and/or coverage. For technical replication, triplicate amplicons of each DNA sample that yielded a PCR product with both primer pairs were sequenced on the Illumina MiSeq platform. Next, we established a computational workflow that allowed us to (i) assess the specificity of the YIHQa and YIHQb PCR assays and (ii) efficiently remove non-*yihQ* sequences from the dataset prior to further diversity analysis (Fig. S2). The workflow consisted of translating nucleic acid sequences into protein sequences, BlastP against the YihQ reference database, and phylogenetic placement in the YihQ reference tree using the evolutionary placement algorithm in RAxML, which shows high accuracy for classification of functional gene sequences [43]. Both primer pairs amplified non-target sequences across all tested samples, but their relative abundance in the sequence libraries varied with sample type (Fig. 2A and B, Table S5). PCR specificity was highest for intestinal samples, particularly for rumen fluid samples, with >98% of reads and >97% of ASVs verified as *yihQ* in both PCR assays. In contrast, less than 35% of reads and 40% of ASVs that were recovered with the YIHQa assay from the Canadian sediment sample were verified as *yihQ*. Hardly any verified *yihQ* reads and ASVs were recovered from sediment using the YIHQb assay. Overall, both YIHQa and YIHQb assays provided high specificity for *yihQ* sequence amplification from intestinal DNA samples but showed considerably reduced reliability with marine sediment DNA.

**Fig. 2.**
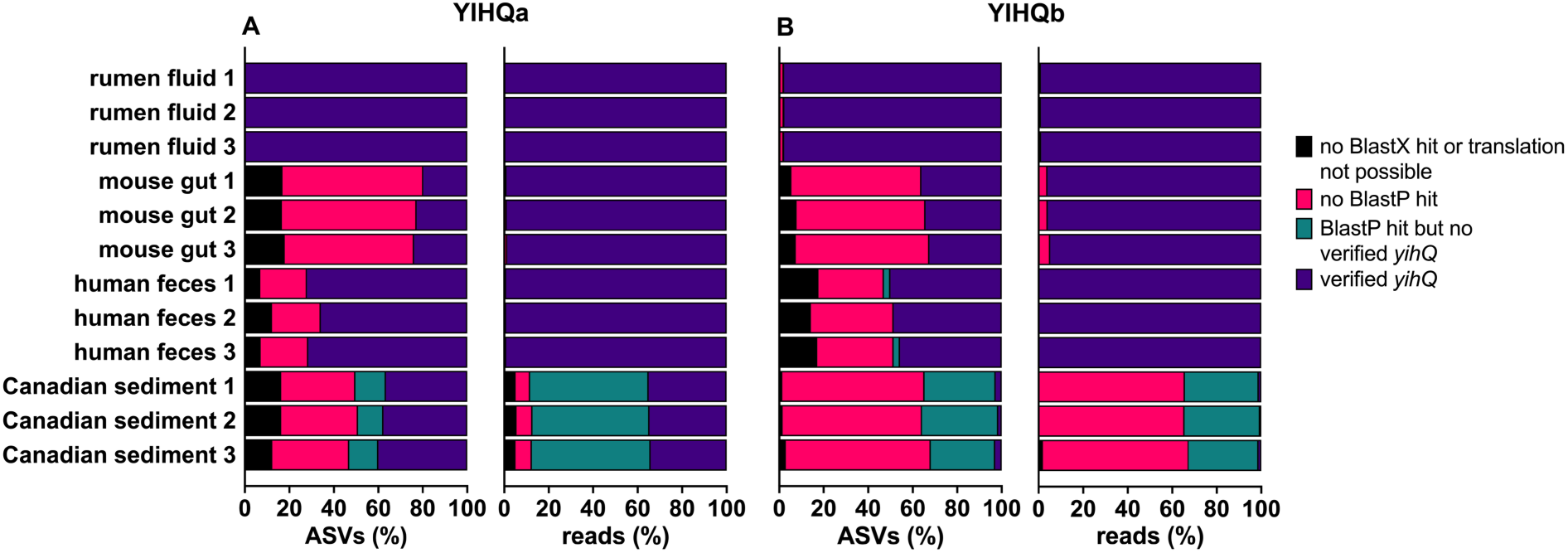
Specificity of the YIHQa and YIHQb amplicon sequencing assays is sample-dependent. Bar plots display the proportion of ASVs or reads (%) recovered from YIHQa (**A**) and YIHQb (**B**) amplicon sequencing assays that were verified as *yihQ* or excluded from further analysis (Fig. S2, Table S5). Each sample was sequenced in triplicates.

Sequences were further clustered into OTUs using a 90% similarity threshold, which has been used for other functional genes, such as *dsrA* and *dsrB* encoding dissimilatory sulfite reductase subunits, for approximate assignment into species-level groups [43]. Alpha diversity analysis at varying library sizes demonstrated high reproducibility among technical triplicates (Fig. S3, Table S6). Rumen fluid samples had the highest *yihQ* richness with more than 1,000 and 400 observed ASVs and OTUs, respectively. Contrarily, mouse gut content and human feces samples, which were pooled from several individuals, had a considerably lower *yihQ* richness with only a few tens of observed ASVs and about ten observed OTUs (Fig. S3). The low species number of potential SQ degraders in the murine and human gut is consistent with prior reports [24,25]. The Canadian sediment sample had an intermediate *yihQ* richness with about 50 and 40 observed ASVs and OTUs, respectively. Accordingly, a library depth of only 100 sequences was sufficient to cover >98% of the expected ASV/OTU richness in the *yihQ* PCR product from the human and mouse gut samples, while hundred times more sequences were required to reach the same coverage for the rumen fluid sample (Fig. S3, Table S6).

The phylogenetic placement of the environmental *yihQ* sequences indicated that both assays recovered similar ASVs that were broadly distributed across the reference tree (Fig. 3). Most ASVs from rumen fluid (64% of 1,220 ASVs from the YIHQa library; 47.5% of 1,548 ASVs from the YIHQb library) were placed at a single, long branch of a YihQ sequence (protein BK011_07555; AUD65555) of *Tenericutes* bacterium MZ-QX. Most ASVs from human feces (66% of 32 ASVs from the YIHQa library; 87% of 15 ASVs from the YIHQb library) and the mouse gut (71% of 17 ASVs from the YIHQa library; 92% of 13 ASVs from the YIHQb library) were affiliated with a *Clostridia* cluster of known SQ degraders, such as *Agathobacter rectalis* [24] and *Otoolea symbiosa* (formerly *Clostridium symbiosum*) [13]. Many rumen-derived ASVs (27% of 1,220 ASVs from the YIHQa library; 37% of 1,548 ASVs from the YIHQb library) were also affiliated with this cluster. Another subset of ASVs from human feces (19% of 32 ASVs from the YIHQa library; 13% of 15 ASVs from the YIHQb library) and the mouse gut (18% of 17 ASVs from the YIHQa library; 8% of 13 ASVs from the YIHQb library) clustered in the *Gammaproteobacteria*, which include the biochemically validated YihQ of *Escherichia coli* [10]. Due to the low coverage of the YIHQb library (only two ASVs were recovered), the phylogenetic placements of ASVs from the two assays could not be compared for the Canadian marine sediment (Fig. 3). However, most marine ASVs from the YIHQa library were placed at a branch with a second YihQ gene copy (protein BK011_09290; AUD64050) of *Tenericutes* bacterium MZ-QX (61% of 71 ASVs) or were affiliated with diverse YihQ sequences within the *Alphaproteobacteria* (27% of 71 ASVs). For refined taxonomic classification, all *yihQ* ASVs generated through the YIHQb assay were additionally matched against GlobDB database, a comprehensive collection of bacterial and archaeal species genomes, using a conservative threshold of 90% YihQ sequence identity (Fig. S4, Table S7). Most ASVs were affiliated with *Oscillospiraceae*, *Lachnospiraceae*, and *Acutalibacteraceae* in the class *Clostridia* (Fig. S4).

**Fig. 3.**
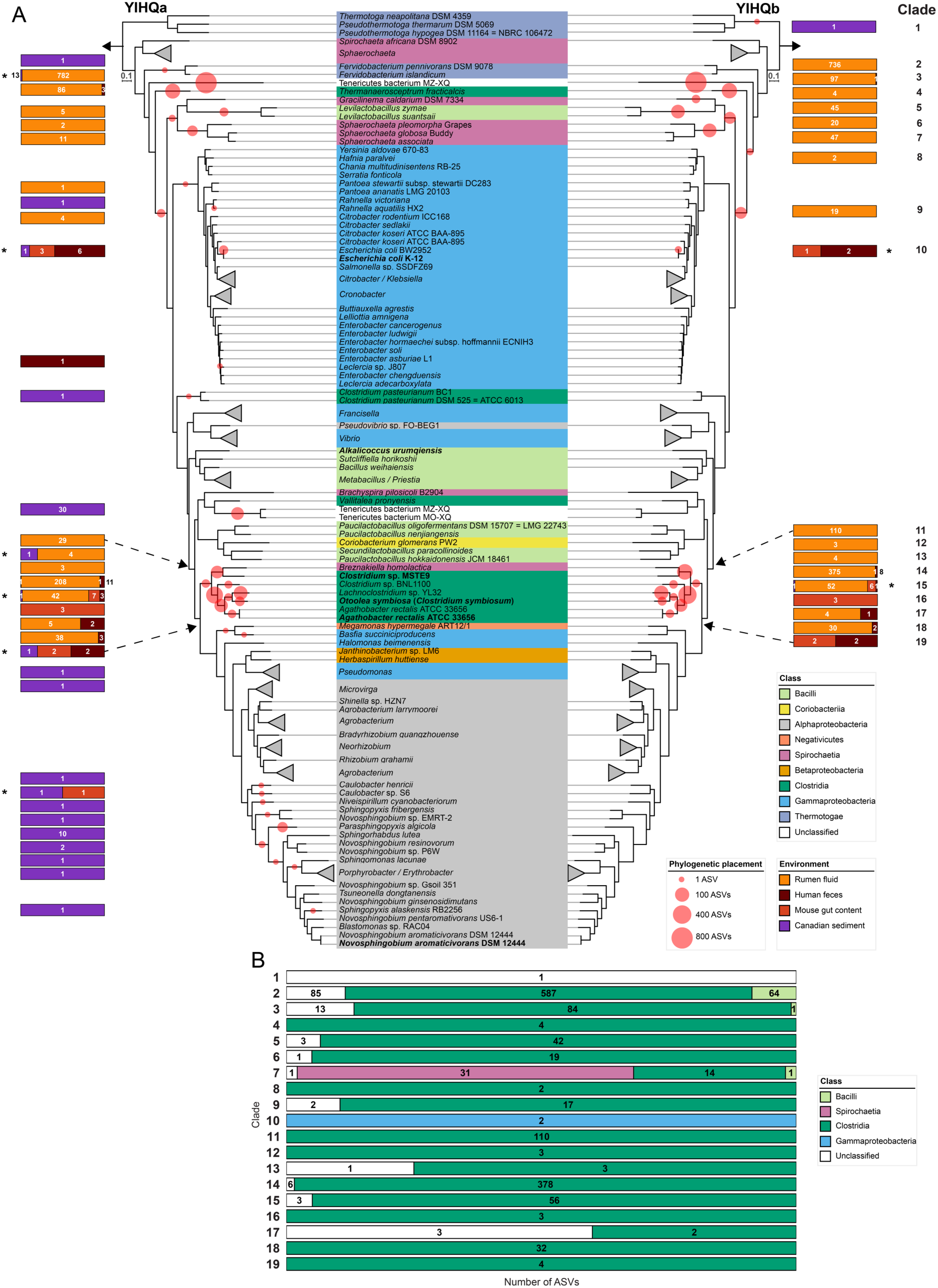
Phylogeny and taxonomic classification of environmental *yihQ* ASVs. Phylogeny of ASVs obtained from human feces, mouse gut content, rumen fluid, and marine sediment using the YIHQa assay (left tree) and the YIHQb assay (right tree) (**A**). For each ASV, a representative, translated sequence was added to the YihQ reference tree using the evolutionary placement algorithm of RAxML. The scale bar shows 10% sequence divergence. The class-level taxonomy of reference sequences is indicated in color. Sequence libraries were not rarefied before placement. Phylogenetic placement is indicated with red circles. The size of each circle represents the number of ASVs placed at that position. Adjacent bars show the percentage distribution of ASVs across environments, with numbers indicating the total ASV count separately for each environment. Asterisks indicate identical ASVs found in different environments. In total, 8 ASVs in YIHQa and 2 ASVs in YIHQb were detected across multiple environments (Table S7). ASV clades in the YIHQb tree are numbered next to the bars. Taxonomic composition of *yihQ* ASV clades generated in the YIHQb amplicon sequencing assay (**B**). Translated *yihQ* ASVs were queried against the GlobDB database using BlastP. The top-scoring hit for each ASV was used for taxonomic assignment at the class-level. ASVs with no match or matches below 90% sequence identity were categorized as unclassified (white). Each horizontal bar represents an ASV clade, as defined in **A**, with individual segments showing the percentage distribution of ASVs to taxonomic classes within that clade. Numbers in bar plots refer to the absolute number of ASVs per class within one ASV clade. Taxonomic assignment of ASVs at the family and genus level is shown in Fig. S4 and Table S7.

The phylogenetic and sequence identity-based analysis provides an initial hypothesis for the taxonomic classification of unknown *yihQ*-containing microorganisms in environmental samples, which must be interpreted with caution. Notably, some organisms, such as *Tenericutes* bacterium MZ-QX and some *Limivicinus* species, possess two copies of *yihQ*. Additionally, the phylogeny of YihQ does not always align with the taxonomy of the organisms (Fig. 3, Fig. S1). For example, numerous *yihQ* ASVs from cow rumen were assigned to the genus *Limivicinus* but form multiple phylogenetically distinct clades in the YihQ tree (Fig. 3, Fig. S4). These potential gene duplications and/or lateral transfers suggest that the evolutionary history of *yihQ* does not necessarily reflect the phylogenetic relationships of the host organisms. Similar to functional marker genes used for other microbial guilds, such as *aprBA*, *dsrAB*, and *amoA* [44–46], this complex evolutionary pattern presents challenges in taxonomic classification but does not preclude the use of *yihQ* as a marker for studying the environmental diversity of SQDG/SQ-degrading microorganisms.

### Novel bacterial species degrade SQ to isethionate via the sulfo-transketolase pathway in cow rumen fluid microcosms

SQDG can be an abundant component of edible green plants and algae [2] and is thus a substrate for the intestinal microbiome. The identity and ecophysiology of intestinal SQDG/SQ-degrading bacteria has so far been only studied for humans [24] and mice [25]. Here, we performed rumen fluid microcosm experiments to investigate microbial SQ degradation in grass- and hay-fed cows, which presumably ingest substantial amounts of SQDG. The SQDG concentration in rumen fluid was 21 µg/g dry weight (Table S8). For comparison, the SQDG concentrations in plants range between 46 µg/g in garlic and 824 µg/g in spinach [47]. SQDG molecules vary in their fatty acid composition, with differences in chain length and degree of saturation. These variations arise because plants produce a diversity of SQDG species depending on their physiology and environmental conditions [48]. In the rumen fluid, the SQDG species 18:3_16:0 represented about 70% of the total SQDG pool, which is similar to the SQDG species composition in green plants such as spinach, basil, and lettuce [49,50].

SQ concentrations decreased only slightly in SQ supplemented rumen fluid microcosms within the first 48 hours of anoxic incubation (Fig. 4A and B). However, SQ was completely metabolized after 168 h. SQ degradation coincided with the production of isethionate while other known organosulfonate SQ degradation products, such as DHPS or 3-sulfolactate [10–13], were not detected. Sulfide concentrations increased across all treatment groups but were significantly higher in SQ-amended microcosms compared to the unamended control (Fig. 4C). Temporal dynamics of acetate, lactate, propionate, and butyrate were not consistently different between SQ-amended microcosms and unamended controls (Fig. S5). However, the production of the known SQ fermentation products acetate and butyrate [12,24,25,29] may have been masked by the high background concentrations of volatile fatty acids (Fig. S5), which is in agreement with previous studies [51,52]. Despite the high richness of *yihQ*-containing microorganisms in the rumen, which is mostly attributed to members of the genus *Limivicinus* (*Oscillospiraceae*) (Fig. S3, Tables S6 and S9), we identified only a few 16S rRNA gene- and *yihQ-*ASVs/OTUs that were significantly enriched in SQ-amended microcosms compared to unamended controls (Tables S10 and S11). 16S rRNA-ASV 1 and 2 were related to the genera *Caproiciproducens* (*Acutalibacteraceae*) and *Mailhella* (*Desulfovibrionaceae*) and reached a maximal relative abundance of about 2.7% and 0.7% in SQ-amended microcosms, respectively (Fig. 4D and E). Additionally, seven *yihQ*-ASVs that clustered in two OTUs were significantly enriched in SQ-amended microcosms. OTU 1 (ASVs 2-7), classified as *Limivicinus* (*Oscillospiraceae*), and OTU 2 (ASV 1), classified as an unknown *Sphaerochaetaceae* bacterium, reached a maximal relative *yihQ* abundance of 12.6% and 9% in SQ-amended microcosms, respectively (Fig. 4F and G, Fig. S6, Table S11).

**Fig. 4.**
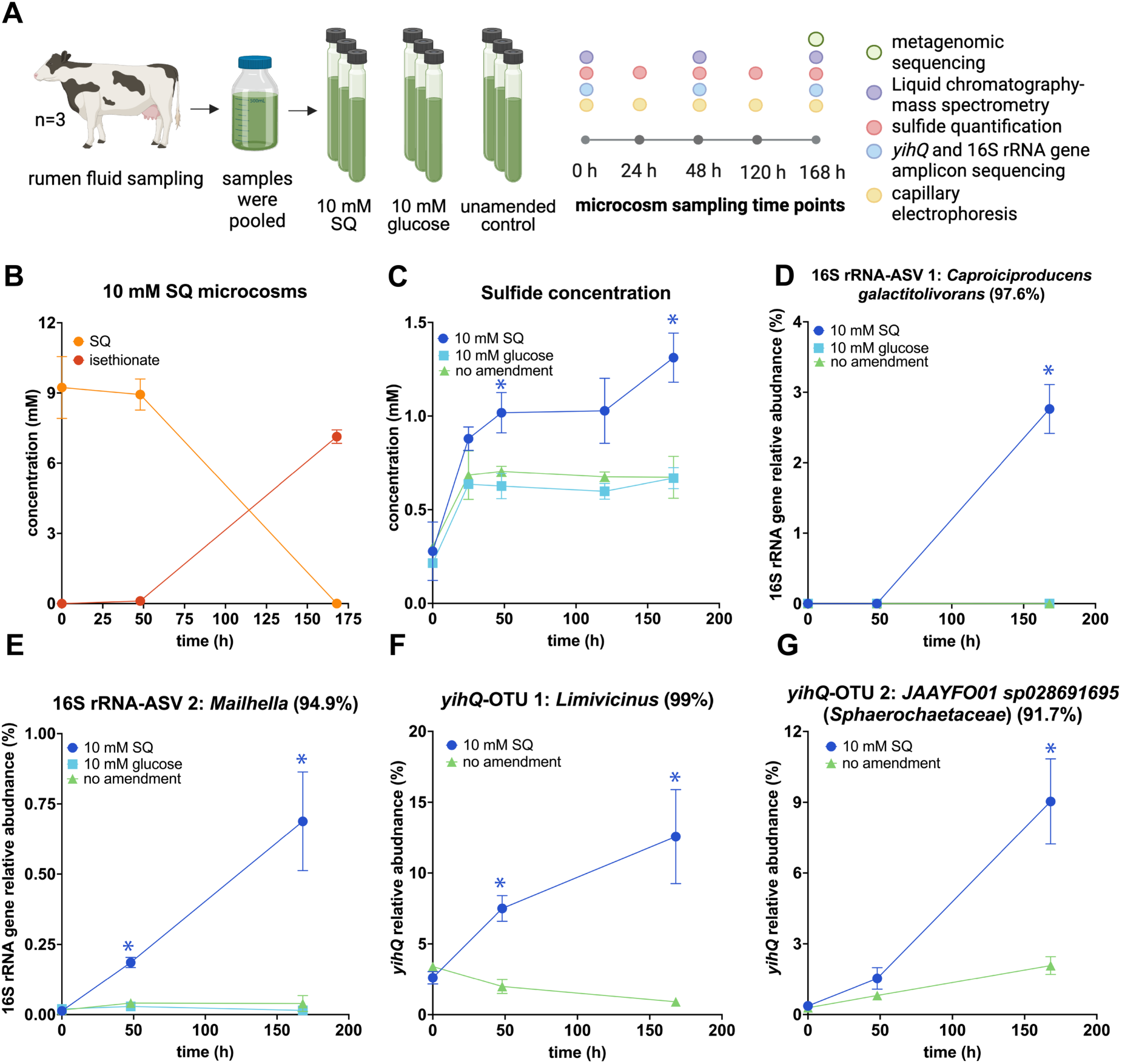
Yet uncultured *Caproiciproducens*, *Limivicinus*, *Mailhella*, and *Sphaerochaetaceae* species are involved in anaerobic degradation of sulfoquinovose to isethionate and sulfide in cow rumen microcosms. Overview of the rumen fluid microcosm experiment and the analytical approach (created with biorender.com) (**A**). Microcosms were amended with 10 mM SQ (n = 3) or 10 mM glucose (n = 3). Unamended microcosms (n = 3) served as controls. SQ (orange) degradation coincided with the production of approx. 7 mM isethionate (red) (**B**). Increase in sulfide concentration was significantly higher in SQ-amended microcosms (dark blue) compared to unamended controls (green) (**C**). Relative abundance dynamics of 16S rRNA gene-ASVs and *yihQ*-OTUs that were significantly enriched in SQ-amended microcosms compared to unamended controls (**D**-**G**). Sequence identity (%) to the next relative is shown in brackets. Data points and error bars represent the average of triplicate measurements and one standard deviation, respectively. Coloured asterisks show significant differences between the respective treatment group and the unamended control at individual time points. Significant differences were determined by ANOVA with Tukey’s posthoc test (**C-E**). Adjusted *p*-values below 0.05 were considered significant. In **G** and **F**, significant differences were determined via DeSeq2 analysis with *p*-values of below 0.01 considered significant. SQ, sulfoquinovose; ASV, amplicon sequencing variant; OTU, operational taxonomic unit.

Accordingly, genome-centric metagenomics recovered 25 medium-to high-quality MAGs from the SQ-amended microcosms, including *Caproiciproducens*, *Limivicinus*, *Sphaerochaetaceae*, and *Mailhella* MAGs (Table S12). The latter MAGs represent yet uncultivated and unnamed species and/or genera. *Caproiciproducens* MAGs SQrumen1 and SQrumen2 each represent a different species (93.3% ANI) [40]. *Caproiciproducens* MAG SQrumen2 belongs to the GTDB species *Caproiciproducens* sp000752215 (GCA_000752215) (97.1% ANI), while *Caproiciproducens* MAG SQrumen1 (93.3% ANI to *Caproiciproducens* sp000752215, GCA_000752215) represent a yet unknown species. The two *Limivicinus* MAGs SQrumen19 and SQrumen23 are only about 50% complete and were thus not used for ANI/AAI calculations. *Limivicinus* MAG SQrumen19 was classified as the GTDB species *Limivicinus* sp900317375 (GCA_900317375), while *Limivicinus* MAG SQrumen23 may represent a yet unknown species (Table S12). *Sphaerochaetaceae* MAG SQrumen3 belongs to the GTDB species JAAYFO01 sp028691695 (GCA_028691695) (99% ANI) and has <53% AAI to species outside the GTDB genus JAAYFO01, suggesting it represents a yet undescribed genus [53]. *Mailhella* MAG SQrumen6 belongs to the GTDB species *Mailhella* sp902783285 (GCA_902783285) (98.8% ANI). Consistent with the metabolite data, the *Caproiciproducens* and *Sphaerochaetaceae* MAGs and *Limivicinus* sp900317375 (GCA_900317375) encode the sulfo-transketolase pathway variant for SQ degradation that produces isethionate (Fig. 5A) [14]. Furthermore, a more comprehensive analysis of these taxa revealed that 10.7% of 28 *Caproiciproducens*, 35.5% of 313 *Limivicinus* (4.7% of 3,230 *Oscillospiraceae*), and 11.8% of 246 *Sphaerochaetaceae* genomes in the GlobDB database have the capacity for SQ degradation via this pathway. The three previously available genomes of the *Sphaerochaetaceae* genus JAAYFO01 do not encode the sulfo-transketolase pathway (Table S13). Notably, all sulfo-transketolase pathway-encoding *Caproiciproducens* genomes lacked genes for the two known sulfoquinovosidases YihQ and SqgA [23] (Fig. 5B). This explains the absence of enriched *yihQ*-ASVs of this genus in the SQ-supplemented rumen microcosms (Table S11) and suggests that SQ-degrading *Caproiciproducens* species either rely on other microorganisms for SQ release or harbor a yet unknown sulfoquinovosidase. Sulfo-transketolase pathway-encoding *Caproiciproducens* genomes additionally contained *sftT*, encoding 6-deoxy-6-sulfo-D-fructose transaldolase. However, they lacked genes for the 3-sulfolactaldehyde reductase (*sftR*) and dehydrogenase (*sftD*), indicating an incomplete sulfo-transaldolase pathway, which is in line with absence of DHPS and 3-sulfolactate in the microcosms [29]. The sulfo-transaldolase and sulfo-Embden-Meyer-Parnas pathways, which are primarily used for SQ fermentation by gut bacteria of humans and laboratory mice [24,25], employ three carbon atoms of SQ for central carbon metabolism and excrete a C_3_-sulfonate (DHPS, 3-sulfolactate) [10,13,14]. In comparison, the sulfo-transketolase pathway of the identified cow rumen bacteria utilizes four SQ carbon atoms for the central carbon metabolism, excretes a C_2_-sulfonate (isethionate), and may thus maximize the energy yield compared to other SQ fermentation pathways [14,54].

**Fig. 5.**
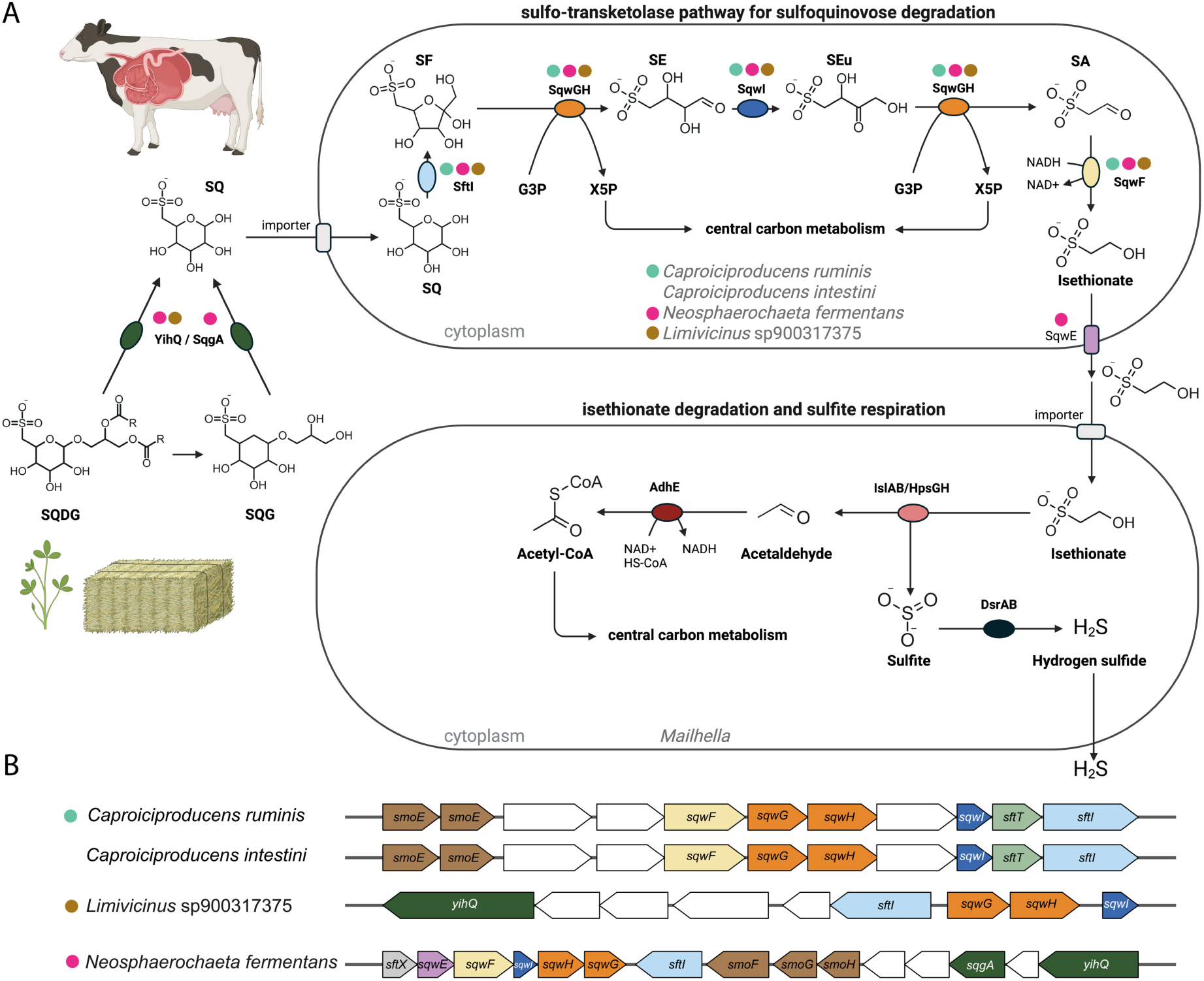
Sulfoquinovose is cooperatively degraded to sulfide via isethionate cross-feeding from diverse sulfo-transketolase-pathway-encoding *Clostridia* species to organosulfur-respiration-encoding *Mailhella* species. (**A**) Cows ingest sulfoquinovosyl diacylglycerol (SQDG) through hay, grass or dietary algae supplements. Newly described bacteria from the cow rumen have the capacity for sulfoquinovose (SQ) catabolism. The sulfoquinovosidases YihQ or SqgA initially cleave SQ from sulfoquinovosyl diacylglycerol (SQDG) or sulfoquinovosyl glycerol (SQG). SQ is further catabolized to isethionate by the sulfo-transketolase pathway-encoding rumen species *Caproiciproducens ruminis* MAG SQrumen1, *Caproiciproducens intestini* MAG SQrumen2, *Neosphaerochaeta fermentans* MAG SQrumen3, and *Limivicinius* sp900317375 (GCA_900317375). Released isethionate is taken up by *Mailhella* species and desulfonated to acetaldehyde and sulfite by the isethionate sulfite-lyase (IslAB) and/or DHPS sulfite-lyase complexes (HpsGH). Sulfite is used as an electron acceptor for anaerobic respiration via the DsrAB-dissimilatory sulfite reductase system. The figure was created with biorender.com. Coloured dots above enzymes indicate presence of the respective gene in the genomes of *Caproiciproducens ruminis* MAG SQrumen1 and *Caproiciproducens intestini* MAG SQrumen2 (turquoise), *Neosphaerochaeta fermentans* MAG SQrumen3 (pink), and *Limivicinus* sp900317375 (brown). Functionally equivalent enzymes are grouped under a single name based on the first characterized homolog (e.g., SftI corresponds to SqvD). (**B**) Gene clusters for SQ degradation via the sulfo-transketolase pathway in *Caproiciproducens intestini*, *Caproiciproducens ruminis*, *Neosphaerochaeta fermentans* and *Limivicinus* sp900317375 (GCA_900317375). Genes depicted in white have no assigned function in sulfur metabolism. YihQ and SqgA, sulfoquinovosidase; SftI, SQ isomerase; SqwGH, 6-deoxy-6-sulfofructose transketolase, SqwI, 4-deoxy-4-sulfoerythrose isomerase; SqwF, sulfoacetaldehyde reductase; SqwE, isethionate exporter; IslAB, isethionate sulfite-lyase; HpsGH, DHPS sulfite-lyase; DsrAB, dissimilatory sulfite reductase; AdhE, aldehyde-alcohol dehydrogenase; SmoE, sulfoquinovosyl glycerol transport ATP-binding protein; SftT, 6-deoxy-6-sulfofructose transaldolase; SmoFGH, transporter; SftX, DUF4867. SQ, sulfoquinovose; SF, 6-deoxy-6-sulfofructose; SE, 4-deoxy-4-sulfoerythrose; SEu, 4-deoxy-4-sulfoerythrulose; G3P, glycerol-3-phosphate; X5P, xylulose-5-phosphate; SA, sulfoacetate; DHPS, 2,3-dihydroxypropane-1-sulfonate.

The slightly increased sulfide production in SQ-amended microcosms likely resulted from anaerobic respiration of isethionate by a *Mailhella* species (Fig. 4C and 5A). The genus *Mailhella* forms a monophyletic group with the genera *Bilophila* and *Taurinivorans*, which comprise important organosulfonate-respiring bacteria from the human and mouse gut, respectively [55]. *Mailhella* MAG SQrumen6 and 75% of 48 *Mailhella* genomes harbour genes that are homologous to the paralogous genes for DHPS sulfite-lyase (*hpsGH*) and isethionate sulfite-lyase (*isIAB*) complexes of *B. wadsworthia* [15,24,56]. Notably, both HpsG and IslA can cleave isethionate to sulfite and acetaldehyde, yet HpsG shows higher activity for DHPS desulfonation [15,56]. Sulfite respiration to sulfide is catalyzed by the dissimilatory sulfite reductase *dsrAB* system, which is evolutionary conserved in *Desulfobacterota* members, including *Mailhella* [55,57].

In summary, our results indicate that diverse, previously undescribed bacterial species in the rumen cooperatively mineralize plant-derived SQ to hydrogen sulfide through isethionate-cross-feeding. These findings carry implications for animal husbandry, as increased sulfite respiration has been associated with decreased activity of methanogenic archaea [58]. Various micro- and macroalgae are tested as feed additives to reduce emissions of the greenhouse gas methane from ruminants [59,60]. Future research may investigate whether SQDG present in algae contributes to their anti-methanogenic properties [61].

In accordance with the SeqCode [62], we propose the following new taxa for three high-quality MAGs (Fig. 5): *Caproiciproducens ruminis* sp.nov. MAG SQrumen1, *Caproiciproducens intestini* sp.nov. MAG SQrumen2, and *Neosphaerochaeta fermentans* gen. nov., sp.nov. MAG SQrumen3.

### Description of Caproiciproducens ruminis sp. nov

*Caproiciproducens ruminis* sp. nov. (ru’mi.nis. L. gen. neut. n. *ruminis*, of the rumen, the source of the bacterium). The designated DNA sequence is MAG SQrumen1 recovered from cow rumen. This species was enriched in anoxic incubations of rumen fluid with sulfoquinovose. Genome analysis suggests sulfoquinovose is fermented to isethionate via the sulfo-transketolase pathway. The NCBI BioSample number for the genome is SAMN48198398.

### Description of Caproiciproducens intestini sp. nov

*Caproiciproducens intestini* sp. nov. (in.tes.ti’ni. L. gen. neut. n. *intestini*, of the gut, referring to the origin of the bacterium). The designated DNA sequence is MAG SQrumen2 recovered from cow rumen. This species was enriched in anoxic incubations of rumen fluid with sulfoquinovose. Genome analysis suggests sulfoquinovose is fermented to isethionate via the sulfo-transketolase pathway. The NCBI BioSample number for the genome is SAMN48198399.

### Description of Neosphaerochaeta fermentans sp. nov

*Neosphaerochaeta fermentans* sp. nov. (fer.men’tans. L. part. adj. *fermentans*, fermenting) The designated DNA sequence is MAG SQrumen3 recovered from cow rumen. This species was enriched in anoxic incubations of rumen fluid with sulfoquinovose. Genome analysis suggests sulfoquinovose is fermented to isethionate via the sulfo-transketolase pathway. The genome additionally encodes two different sulfoquinovosidases (YihQ and SqgA). The NCBI BioSample number for the genome is SAMN48198401.

### Description of *Neosphaerochaeta* gen. nov

*Neosphaerochaeta* gen. nov. (Ne.o.sphae.ro.chae’ta. Gr. masc. adj. neos, new; N.L. fem. n. *Sphaerochaeta*, a bacterial genus; N.L. fem. n. *Neosphaerochaeta*, a new genus related to *Sphaerochaeta*). Type species: *Neosphaerochaeta fermentans* sp. nov., family: *Sphaerochaetaceae*.

## Conclusions

We developed the first functional gene amplicon sequencing assays for analyzing the diversity of SQDG/SQ-degrading microorganisms in complex samples. The two PCR assays both target the same gene region of the YihQ sulfoquinovosidase and exhibited similar environment-dependent specificity. They were highly specific for *yihQ* in intestinal samples from humans, mice, and cows but amplified a substantial proportion of non-*yihQ* sequences from marine sediments. To ensure accurate *yihQ* diversity analysis, we implemented a computational workflow to efficiently remove non-target sequences. However, the potential amplification of non-target genes may reduce the sensitivity of the PCR assays in certain environments, such as marine sediments. Future improvements to the *yihQ*-targeted PCR assays could address these limitations. Additionally, for a more comprehensive assessment of SQDG/SQ-degrading microorganisms, functional gene amplicon sequencing should be complemented by PCR assays targeting the more recently identified SqgA sulfoquinovosidase [23].

Among all analyzed samples, rumen fluid exhibited the highest *yihQ* richness, suggesting that the species/strain-level diversity of potential SQDG/SQ-degraders in the cow gut is one to two orders of magnitude greater than in the human and mouse gut. Furthermore, yet uncultured species were enriched in rumen fluid microcosms supplemented with SQ and identified as primary (*Caproiciproducens*, *Limivicinus*, *Neosphaerochaeta*) or secondary (*Mailhella*) SQDG/SQ-degraders through metabolite analysis, amplicon sequencing, and comparative genomics. This study revealed SQDG/SQ degradation as a relevant metabolic trait of the cow rumen microbiome and provided insights into the diversity and metabolic potential of the newly described taxa involved.

## Acknowledgments

This research was funded in part by the Austrian Science Fund (FWF) [grant DOI <u>10.55776/DOC69</u>, <u>10.55776/P31996</u>, and <u>10.55776/COE7</u>] and the EU MSCA postdoctoral fellowships to S.-C.C. (action number 101059607, DOI <u>10.3030/101059607</u>, DatingSuCy) and T.S.T. (action number 101205556, DOI <u>10.3030/101205556</u>, SLIDES). For open access purposes, the authors have applied for a CC BY public copyright license to any author-accepted manuscript version arising from this submission. We thank Cameron Strachan for sampling of rumen fluid, Anna Müller and Jillian Petersen for sampling of the marine sediments, and Bernhard Schink for help with Latin naming of taxa.

## Author contributions

J.K. and A.L. conceived the study, with support from M.M.. J.K., H.S., P.M., S.B., and R.S. performed experiments and analyzed data. D.R.S., E.S., W.W., and D.S. contributed essential samples or data, experimental infrastructure, analytical instrumentation and/or expert advice. J.K., S.C., T.S.T., J.O., J.S., and B.H. performed bioinformatic analyses. J.K., S.C., P.M, S.B., R.S., M.M., and A.L. interpreted the data. J.K. and A.L. wrote the article. All authors revised and approved the manuscript.

