## Supplementary Materials & Methods and Figures 1-6 for "Extensive richness and novel taxa of sulfoquinovose-degrading bacteria in the cow rumen"

### Supplementary Materials and Methods

#### Design of *yihQ*-targeted PCR assays

A YihQ reference database was created using sequences of biochemically validated YihQ sulfoquinovosidases and closely-related homologs of functionally different enzymes, such as alpha-xylosidases, of the glycoside hydrolase family 31 as queries for Blast against the KEGG prokaryotes database [1] (Table S1). The retrieved YihQ homologs (n = 500) were de-replicated at 95% protein sequence identity, and the resulting sequences (n = 294) were further aligned with the biochemically validated proteins using MUSCLE [2]. A maximum-likelihood tree was constructed using FastTree [3] with a gamma distribution (-gtr option) to account for different evolution rates at different sites and the Whelan and Goldman protein evolution model (-wag option) [4]. Bootstrap support values for each tree branch was calculated using the CompareToBootstrap.pl script [5]. Homologs (n = 131) that formed a well-supported (>70% bootstrap support) monophyletic clade with biochemically validated YihQ sequences in the tree served as the basis for primer design (Fig. S1). After manually identifying conserved sites in the YihQ sequence alignment, the corresponding sites in the nucleic acids sequence alignment were used for the design of degenerate primers. To further minimize nucleotide degeneracy for some primers, the alignment was examined for each degenerate nucleotide in the primer sequence to determine whether a less degenerate alternative could be used.

Genomic DNA extracts from the *yihQ*-encoding strains *Escherichia coli* K12, *Agathobacter rectalis* A1-86, and *Enterocloster clostridioformis* YL32 were used as positive controls for PCR. DNA of *Segatella copri* DSM 18205 and ultrapure water served as non-target control and negative control, respectively. All strains were grown anaerobically (85% $N_2$, 10% $CO_2$, 5% $H_2$)

in brain heart infusion broth at 37°C until the stationary phase, after which genomic DNA was extracted using Chelex 100 Resin (Bio-Rad, Austria), as previously described [6]. Briefly, 200 µl of ultrapure water was added to 200 µl liquid bacterial culture and centrifuged at 15,500 g for 2 min. The supernatant was discarded. 100 µl of 5% Chelex 100 resin was added to the pellet and contents were vortexed until homogenized. Samples were incubated at 100°C for 10 min, then again centrifuged at 15,500 g for 2 min and DNA-containing supernatants were transferred to a new tube. DNA concentration was determined using a QubitTM 3 Fluorometer (Invitrogen, USA). DNA extracts were stored at -20°C using them for PCR. The absence of PCR-inhibiting substances in the DNA extracts was verified by a positive 16S rRNA gene-targeted PCR with the universal primers 515F (5'-GTG YCA GCM GCC GCG GTA A) and 806R (5'-GGA CTA CNV GGG TWT CTA AT) [7,8].

All *yihQ*-targeted primer pairs were first tested using a temperature gradient PCR. The initial PCR program consisted of denaturation at 95°C for 3 min, followed by 30 cycles of 95°C for 30 s, annealing at temperatures ranging from 40°C to 65°C in ~1°C increments for 30 s, and elongation at 72°C for 45 s, with a final elongation step at 72°C for 10 min. Each 50 µl reaction mixture contained 1× DreamTaq buffer, 0.2 mM dNTPs, 1.25 U recombinant Taq polymerase (5 U/µl), 10 µg bovine serum albumin, 10 µM of each primer, and 1 µl of template DNA. All reagents were sourced from Thermo Fisher Scientific (Germany).

Primer pairs that yielded PCR products of the expected size with positive controls and showed no amplification in negative or non-target controls were validated via Sanger sequencing. To optimize assay conditions, primers that successfully amplified the target gene were further tested using touchdown PCR. Annealing temperatures of 65–55°C, 64–54°C, 63–53°C, 62–52°C, 61–51°C, and 60–50°C were tested. In each condition, the annealing temperature started at the highest value of the respective range (e.g., 65°C for 65–55°C) and decreased by

1°C per cycle over the first 10 cycles. After reaching the lowest temperature of the range (e.g., 55°C for 65–55°C), the reaction continued for an additional 20 cycles at this final annealing temperature. Then, using the determined optimal touchdown temperature range, magnesium chloride concentrations between 0.5–7 mM were tested in 0.5 mM increments. Successful amplification of target genes was confirmed via Sanger sequencing.

The final PCR program was tested with DNA from several environmental and intestinal samples and included an initial denaturation at 95°C for 3 min, followed by 10 cycles of 95°C for 30 s, 63–53°C for 45 s (decreasing 1°C per cycle), and 72°C for 45 s. This was followed by 20 cycles of 95°C for 30 s, 53°C for 45 s, and 72°C for 45 s, with a final elongation at 72°C for 15 min. Each 50 µl reaction contained 1× Taq buffer (with potassium chloride), 0.2 mM dNTPs, 1.25 U Taq Polymerase (recombinant, 5 U/µl), 4 mM magnesium chloride, 10 µM of each primer, and 1 µl of template DNA (Table S2). All reagents were obtained from Thermo Fisher Scientific (Germany). PCRs with environmental DNA included DNA from a *yihQ*-encoding strain as positive control and DNA from *S. copri* without *yihQ* and ultrapure water as negative controls.

#### Rumen fluid microcosms

Rumen fluid microcosms were prepared in an anaerobic tent (Coy Laboratory Products, USA) operated with a gas mix of 85% $N_2$, 10% $CO_2$, and 5% $H_2$, maintaining $H_2$ levels of 1.5–2.5%. Rumen fluid samples from three cows were pooled (1:1:1 ratio, v/v/v ) and homogenized. Large particles were left to settle. In each Hungate tube, rumen fluid was mixed 1:1 with 1× PBS (pH 7.4), including the specific amendments, to a final volume of 10 ml. Microcosms were amended with 10 mM SQ (MCAT GmbH, Germany), 10 mM D-glucose (Carl Roth, Germany) or left unamended. Triplicate microcosms were set up for each condition. Tubes were sealed with butyl-rubber stoppers and screw caps. Incubations were conducted at 37°C, with sampling over seven days. Sampling for DNA extraction and metabolite analysis was done after 0, 24, 48, 120, and 168 h. Microcosms were subsampled by retrieving one ml aliquots of fluid, which were centrifuged at 11,000 g for 10 min at 4°C. Pellets were stored at −80°C for DNA extraction. Supernatants were stored at −20°C for metabolite analyses.

#### Metabolite analysis

Sulfide was quantified in subsamples of 20 µl culture that were fixed by adding 2 g/l zinc acetate in the anaerobic tent. Fixed samples were measured colorimetrically using the Infinite M Nano+ microplate reader (Tecan, Austria), as previously described [9]. Due to the high amount of particles in the rumen fluid microcosms, sampling with needle and syringe was not possible and thus rubber stoppers were briefly removed for sampling. Due to possible leakage, sulfide concentrations might therefore be underestimated in all treatments and time points.

Formate, acetate, propionate, butyrate, valerate, and lactate were quantified in rumen fluid microcosms using an Agilent 7100 Capillary Electrophoresis system (Agilent, USA). Culture supernatants were diluted 1:5 to 1:20 in ultrapure water containing 0.1 mM succinate as an internal standard. Calibration standards for each metabolite were prepared at concentrations of 0.025, 0.05, 0.1, 0.25, 0.5, and 1 mM. An organic acid buffer (Agilent, USA) was used as the running buffer. The bare fused silica capillary (75 µm inner diameter, 72 cm effective length; Agilent, USA) underwent a 4-minute preconditioning flush before each measurement. After preconditioning, samples were injected at 5 kPa for 5 seconds. Electrophoretic separation was performed at −25 kV with a current of 100 µA and a power of 6 W.

Liquid chromatography–mass spectrometry was used to quantify SQ and its metabolites isethionate, 2,3-dihydroxypropane-1-sulfonate (DHPS), 3-hydroxypropane-1-sulfonate, 3-sulfolactate, and 3-sulfopropionate in supernatants of rumen fluid microcosms. Samples were quantified using a Shimadzu LCMS-2020 single quadrupole mass spectrometer with electron spray ionisation. Separation of organosulfonates was achieved using an iHILIC-Fusion(+) column (100 × 2.1 mm, 3.5 µm, 100 Å), as described previously [10].

Ultra-high-performance reverse phase liquid chromatography combined with mass spectrometry was used to quantify and identify all molecular species of SQDG in rumen fluid. A rumen fluid sample from one cow was subdivided into two replicates (each 0.1 g) (stored at -80°C) and freeze-dried. Lipid extraction was performed as previously reported [11,12], with few modifications. Briefly, six ml of chloroform-methanol-citrate buffer (0.15 M, pH 4.0, 1:2:0.8 v/v/v) was added to freeze-dried samples and mixed in glass vials. Internal standard (1 µg/ml, Equisplash; $^{2}H$-labelled multilipid standard from Avanti Research, USA) was added, vortexed, sonicated for 10 min at room temperature, and incubated overnight in the dark at room temperature. Subsequently, samples were centrifuged (1,500 g for 10 min) and the supernatant was carefully transferred to a clean vial. The pellet was re-extracted with 2.6 ml of the chloroform-methanol-citrate buffer, vortexed, and centrifuged. The supernatants of both extractions were combined. Then, 1.5 ml of chloroform and 1.5 ml each of citrate buffer were added to the combined supernatants, vortexed, and incubated overnight for phase separation. The lower, lipid-rich organic phase was cautiously transferred to pre-weighed vials. Subsequently, the lipid fraction was dried with $N_2$ gas and then solubilized in 200 µl of injection solvent (water:acetonitrile:isopropyl alcohol, 1:2:1, v/v/v). For liquid chromatography, an Ultimate 3000 Ultra-High-Performance Liquid Chromatography system (Thermo Fisher Scientific, Germany) with an Accucore C30 column (150 × 2.1 mm; Thermo Fisher Scientific, Germany) and fused-core particles of 2.6 µm diameter were used. The column oven and autosampler were set at 10°C and 40°C, respectively. Solvent A consisted of acetonitrile and water (3:2, v/v) and solvent B of isopropyl alcohol and acetonitrile (9:1, v/v), each amended with 0.1% formic acid and 10 mM ammonium formate. The flow rate was set to 325 µl/ min with 5 µl injection volume. The gradient profile was as follows: 0-18 minutes, nonlinear increase from 30% to 85% solvent B; 18-20 minutes, ramp to 90% solvent B; 20-24 minutes, hold at 90% solvent B; and 24-28 minutes, linear decrease to 30% solvent B, followed by 4 minutes of re-equilibration at 30% solvent B. Subsequent mass spectrometry was performed with a Q Exactive Orbitrap Mass Spectrometer (Thermo Fisher Scientific, USA) in positive and negative ion modes using a heated electrospray ionisation source. For both polarities, sheath and sweep gas were 40 and 10 (arbitrary units), respectively, and the rate for auxiliary gas was 5 for positive and 7 for negative ionisation modes. The spray voltage was at 3 kV, capillary temperature was at 370°C, the ion transfer tube temperature was at 285°C, for each of the ionisation settings, and the S-lens radio frequency level was adjusted to 50. In separate runs of each sample electrospray ionization-positive and electrospray ionization-negative modes with/without higher-energy collisional dissociation (HCD) fragmentation were measured. The Orbitrap mass analyzer was run at a mass resolving power of 70,000 in positive/negative mode, scan range of 300–1800 m/z, and automatic gain control target of $10^6$. In the HCD mode, the resolving power was set to 17,000, employing HCD fragmentation with stepped normalized collision energy of 20, 25, and 30% (HCD injection time: 100 ms; isolation window: 1 m/z; automatic gain control target: $10^6$), with dynamic exclusion setting of 6 s in the Top5 data-dependent MS2 mode. Raw data were analyzed using the Lipidsearch 5.0 software (Thermo Fisher scientific, USA). SQDG (18:3_16:0) (Sigma-Aldrich, USA) was used as an external standard and class-specific lipid standard for all SQDG-class compounds to quantify SQDG. The Skyline software was used for the identification and quantification of SQDG compounds in the mass spectrometry analysis [13,14].

#### 16S rRNA gene and *yihQ* amplicon sequencing and analyses

Genomic DNA from environmental, intestinal, and rumen fluid microcosm samples was isolated using the DNeasy PowerSoil Pro Kit (Qiagen, Austria). Samples were processed in two ml Lysing Matrix E tubes (MP Biomedicals™, Austria) with CD1 lysis buffer. Samples were incubated at 65°C for 10 minutes and then homogenized using the FastPrep-24 Classic bead-beating system (6 m/s, 2 × 40 seconds; MP Biomedicals™, Austria). DNA was further isolated according to the manufacturer's standard protocol. The concentration of DNA was determined using a QubitTM 3 Fluorometer (Invitrogen, USA). DNA samples were stored at -20°C. The absence of PCR-inhibiting substances in the DNA extracts was verified by a positive 16S rRNA gene-targeted PCR with the universal primers 515F and 806R.

Amplicon sequencing was conducted by the Joint Microbiome Facility (University of Vienna, Medical University of Vienna) under project IDs JMF-2310-13 and JMF-2302-02. DNA from each environmental sample was sequenced in triplicates within the same sequencing run. A two-step PCR method [15,16] and primer pairs YIHQa (YIHQ1201Fa/YIHQ1526Ra) and YIHQb (YIHQ1201Fb/YIHQ1526Rb) were used to amplify and barcode an approximately 350 bp region of the *yihQ* gene (Fig. 1). In the first step, amplification was performed with *yihQ*-targeted primers containing 16 bp 5' head adapters (forward: 5'-GCT-ATG-CGC-GAG-CTG-C-

3′; reverse: 5′-TAG-CGC-ACA-CCT-GGT-A-3′) and with an optimized magnesium chloride concentration of 3 mM. Other reaction components and PCR conditions were as described before (Table S2). In the second step, the individual PCR products were tagged with unique 12 bp barcodes [16].

Samples from the rumen fluid microcosm experiment were analyzed by *yihQ* amplicon sequencing with primer pair YIHQb according to the described protocol and by 16S rRNA gene amplicon sequencing. For 16S rRNA gene amplicon sequencing, the two-step PCR-barcoding method (H_515F_mod-5′-(head) GCT ATG CGC GAG CTG C - GTG YCA GCM GCC GCG GTA A and H_806R_mod-5′-(head) TAG CGC ACA CCT GGT A - GGA CTA CNV GGG TWT CTA AT) was used to target bacteria and archaea [15,16].

Barcoded *yihQ* and 16S rRNA gene amplicon libraries were normalized with the SequalPrep normalisation plate kit and subsequently pooled (TrueSeq Nano Kit, Illumina, USA; modified according to [15]), prior to preparing the sequencing libraries. Samples were sequenced on the MiSeq (V3, 600 cycles, Illumina, USA) and results were analyzed according to the procedures described before [16].

Amplicon sequence variants (ASVs) of 16S rRNA gene sequences were inferred and classified using DADA2 [17] with Release 138 of the SILVA 16S rRNA sequence database [18]. All *yihQ* amplicon sequences were translated into amino acid sequences (Fig. S2). To achieve this, open reading frames were identified and translated using EMBOSS Transeq [19]. Sequences that generated a stop codon in this approach were instead translated with Prodigal in metagenomic mode, which is suitable for partial sequences that lack start codons [20]. If neither method yielded a stop codon-free translation, a local BlastX search was performed against the YihQ reference database [21]. Sequences with a significant BlastX hit (defined as having a coverage >80% and e-value <$10^{-4}$) were translated based on the reading frame of their best BlastX match. Sequences that still retained a stop codon after translation in the suggested BlastX reading frame or lacked a significant BlastX hit in the YihQ reference database were excluded from further analyses. Translated sequences without stop codons from all approaches were compiled and subjected to a BlastP search against the YihQ reference database to ensure that only sequences with significant similarity to sequences in the YihQ reference database were included in the subsequent phylogenetic analysis [21]. Sequences without a significant BlastP hit (coverage >80% and e-value <$10^{-4}$) in the YihQ reference database were excluded from further analyses.

All amino acid sequences with a significant BlastP hit in the YihQ reference database were placed into the YihQ reference tree using the evolutionary placement algorithm of RAxML. Phylogenetic placement was performed with the LG substitution model and a gamma distribution to account for rate heterogeneity across sites [22–24]. Sequences that were placed within the YihQ branch of the reference tree with an accumulated likelihood of >0.95 were classified as verified *yihQ* sequences. ASV sequences that did not meet these criteria were considered non-target sequences, removed from the dataset, and excluded from further analysis. All verified *yihQ* ASV sequences were clustered into operational taxonomic units (OTUs) using vsearch [25] with a 90% sequence identity threshold. Translated verified *yihQ* ASV sequences generated with the YIHQb amplicon sequencing assay were additionally subjected to BlastP analysis against all YihQ sequences extracted from the GlobDB database (version r220) for taxonomic classification. GlobDB is a curated, dereplicated collection of representatives genomes of all species in the global taxonomy database (GTDB) and of species that are not present in GTDB but in the genomic catalog of earth's microbiomes, the searchable planetary-scale microbiome resource, and the genomic catalog of soil microbiomes [26].

Verified *yihQ* sequences were further analyzed in R Studio (rstudio.com, R version 4.2.2) using the software packages “Phyloseq” (v1.42.0) [27] and “Vegan” (v2.6-4) [28]. Libraries were subsampled at 100, 1000, and 10000 reads to evaluate the impact of sequencing depth on diversity estimates. Alpha diversity was evaluated using the observed number of ASVs and OTUs, Shannon diversity index, and Simpson diversity index.

Differential abundance analysis of 16S rRNA gene sequences and verified *yihQ* sequences from the rumen fluid microcosm experiment was conducted using DESeq2 (v1.38.3) in RStudio, as described previously [29]. To mitigate biases from uneven library sizes, sequencing libraries were rarefied to match the smallest library before DESeq2 analysis. For 16S rRNA gene data, SQ-amended microcosms were compared to unamended controls at 48 h and 168 h of incubation. Relative abundances of *yihQ*-ASVs and OTUs were compared between SQ-amended microcosms and controls at 168 h of incubation. Adjusted *p*-values <0.01 were considered significant. Taxonomic classification of enriched 16S rRNA gene ASVs was verified using BlastN against the NCBI 16S rRNA sequence database, excluding sequences from uncultured organisms and environmental samples [30]. The next related YihQ-encoding genome in the GlobDB database of each significantly enriched *yihQ* OTU and ASV was identified by using BlastP.

#### Metagenomics and genome analysis

Metagenomic sequencing was performed with samples from SQ-amended (n = 2) and unamended control (n = 1) microcosms of rumen fluid, taken after 168 h of incubation. Libraries were prepared using the NEBNext Ultra FS II DNA Library Prep Kit (New England Biolabs, USA) for Illumina and sequenced on a NovaSeq6000 (1/2 SP flow cell, 2×100 bp reads; Illumina, USA). Quality control was conducted using a Snakemake v8.20.3 workflow to trim reads based on quality scores [31]. Reads were quality-checked with FastQC v0.12.1 [32], and summary statistics were compiled using MultiQC v1.21 [33]. Adapter sequences and phiX contamination were removed with BBDuk (BBMap v39.06) [34], retaining reads of at least 50 bp. Right-end k-trimming was applied (k-mer = 21, minimum k-mer = 11, hamming distance = 2) with "tpe" and "tbo" options, and quality trimming was performed with a Q-score of 15. The quality-controlled read library was quality checked using FastQC v0.12.1 and quality statistics were merged using MultiQC v1.21 for comparison to the non-trimmed quality. BBMap's reformat.sh was used to interleave the read library.

Reads were assembled with SPAdes v4.0.0 in "meta" mode, using k-mers from 21 to 121 (step size: 10) [35]. Contigs shorter than 1,000 bp were removed using reformat.sh of BBMap. Metabat v2.15 [36] was used to bin putative metagenome-assembled genomes (MAGs) without coverage data (minimum length: 1,500 bp). Coverage information for the assembly was generated with BBMap (≥98% identity) using the read library, followed by sorting with SAMtools v1.20 [37]. A depth file was generated using jgi_summarize_bam_contig_depths and the assemblies were binned into putative MAGs again using the depth file in Metabat. Dereplication of putative MAGs was performed with dRep v3.5.0 [38], applying a minimum completeness threshold of 50% and a maximum contamination threshold of 10%. Taxonomic classification of dereplicated MAGs was conducted using GTDB-Tk v2.4.0's classify_wf (database r220, ANI screening skipped) [39]. Barrnap v0.9 was used to identify and recover 5S, 16S, and 23S rRNA gene sequences in the genomes [40]. Coverage and the percentage of reads mapped to each MAG were determined with BBMap with a mapping identity of 95%. Average amino acid identity (AAI) was calculated using the Enveomics Collection [41]. Whole-genome average nucleotide identity (ANI) was calculated using FastANI (version 1.33) [42].

New Hidden Markov Models (HMMs) were generated for the detection of enzymes and transport proteins of sulfoquinovose, isethionate, and DHPS degradation pathways (Table S3). Initially, the sequences of the biochemically and/or physiologically characterized enzymes and representative sequences of putative transport proteins were used as a query for a Diamond BlastP against the GlobDB (version r220) with a bitscore cutoff of 100. If a protein sequence in GlobDB had hits above the cutoff to more than one query sequence, the query with the highest bitscore was selected and the target sequence in GlobDB was labeled with the name of the query. For all hits in the resulting list of labeled GlobDB sequences the genomic vicinity was analyzed for additional hits to other query sequences. Specifically, for each genome the nucleotide distance between all genes was calculated. Genes with a maximum of 3500 nucleotides of intergenic space were considered to form a gene cluster, if at least five genes were encoded and if these were homologous to five different query sequences. An exception was made for the isethionate sulfite-lyase IslA, DHPS-sulfite-lyase HpsG, and DHPS-dehydratase HpfG, as these occur in gene clusters of less than five genes [43,44]. Here, only gene cluster arrangements that were specifically described in the respective publication were selected. Across all genomes in GlobDB, the gene clusters encoding the same number and types of homologs were then grouped into unique gene cluster groups. In the next step, a gene cluster group was excluded if it did not contain at least one gene cluster with one encoded protein with 90% identity to the respective query sequence. To generate the final sequence set for each functionally different protein, all homologs encoded in any gene cluster group were merged into an individual sequence collection for each initial reference protein. The protein sequences in each collection were then aligned and used to generate a HMM, with trusted and noise cutoff scores as previously described [45]. HMMs and cutoff scores were integrated into HMSS2 [46], which was then used to identify sulfur metabolism genes in the recovered MAGs and genomes from the families *Sphaerochaetaceae* and *Oscillospiraceae*, the genera *Mailhella*, *Caproiciproducens* and JAAYFO01 in GlobDB. Proteins were only counted as present (i) if they were detected with a bitscore above the respective trusted cutoff or (ii) if they were detected with a bitscore above the noise cutoff and had at least 95% identity to one of the sequences used to generate the HMM.

### Supplementary Figures

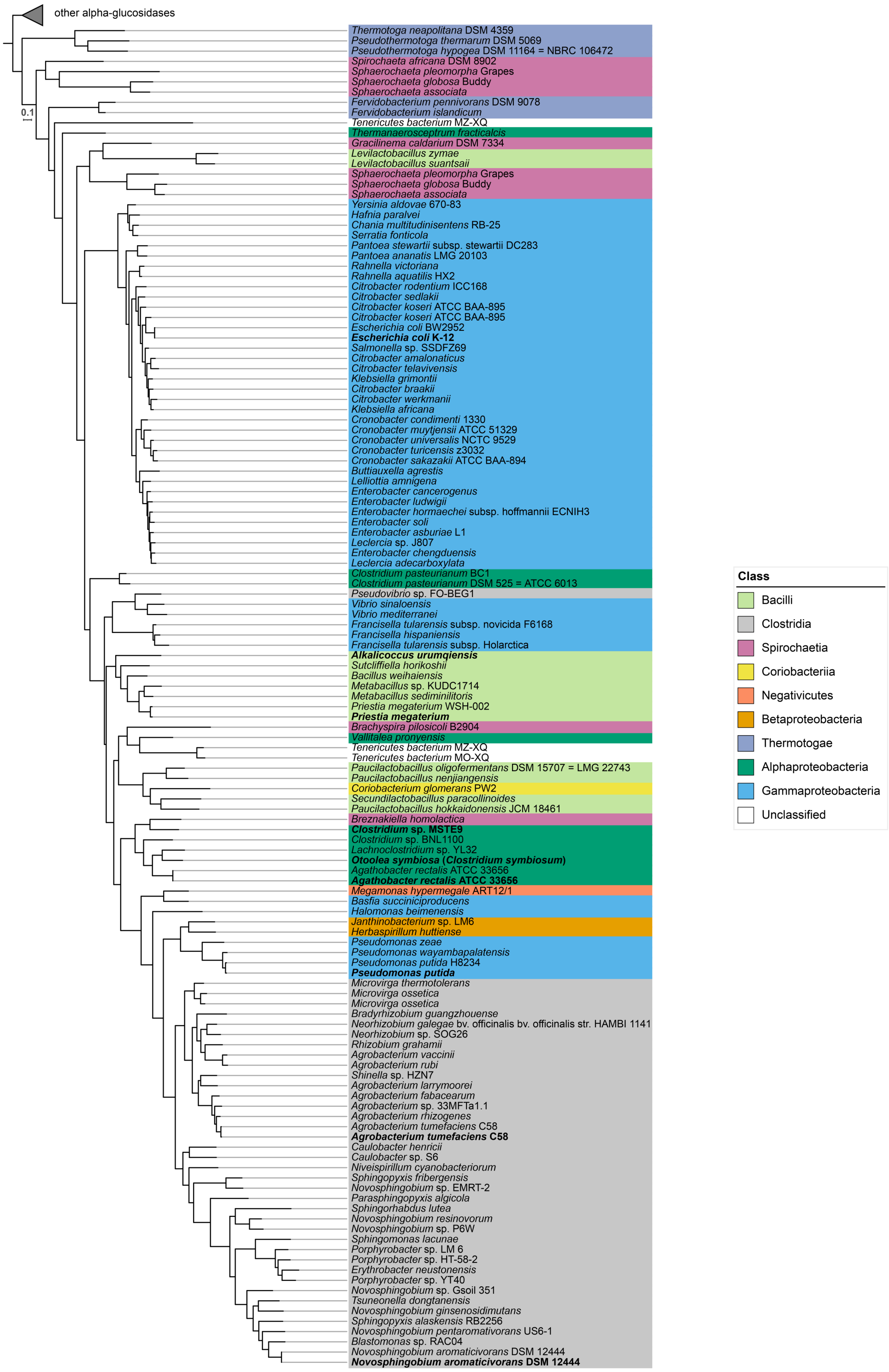

**Figure S1. Phylogeny of YihQ sulfoquinovosidases.**

The maximum likelihood YihQ reference tree was calculated with 131 YihQ sequences and 163 outgroup sequences from the reference database after de-replication at 95% protein sequence identity. Outgroup sequences are collapsed. Biochemically or physiologically verified YihQ sulfoquinovosidases are highlighted in bold. Colors indicate taxonomic assignment at the class level.

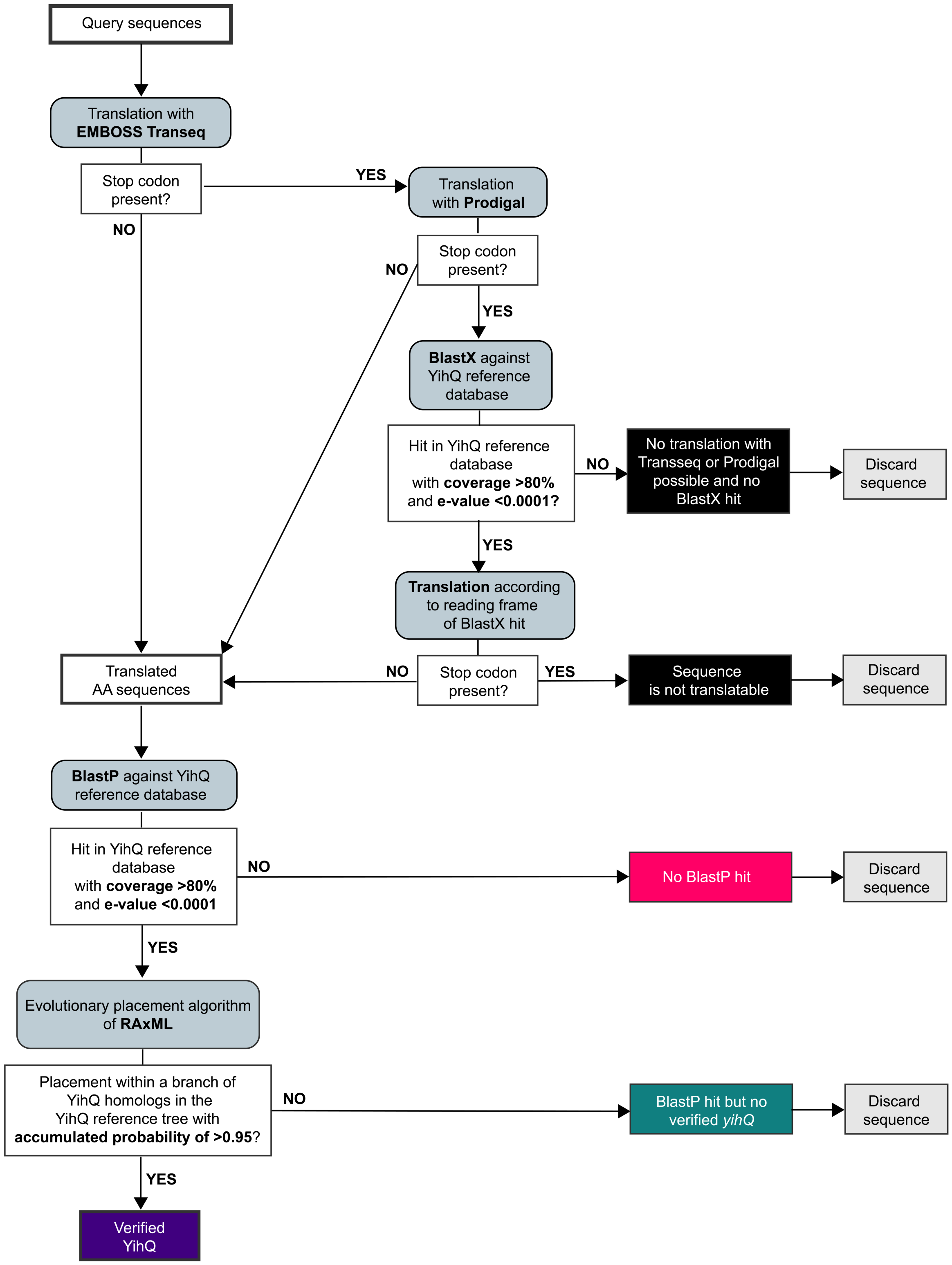

**Figure S2. Computational workflow for identification and filtering of *yihQ* sequences.**

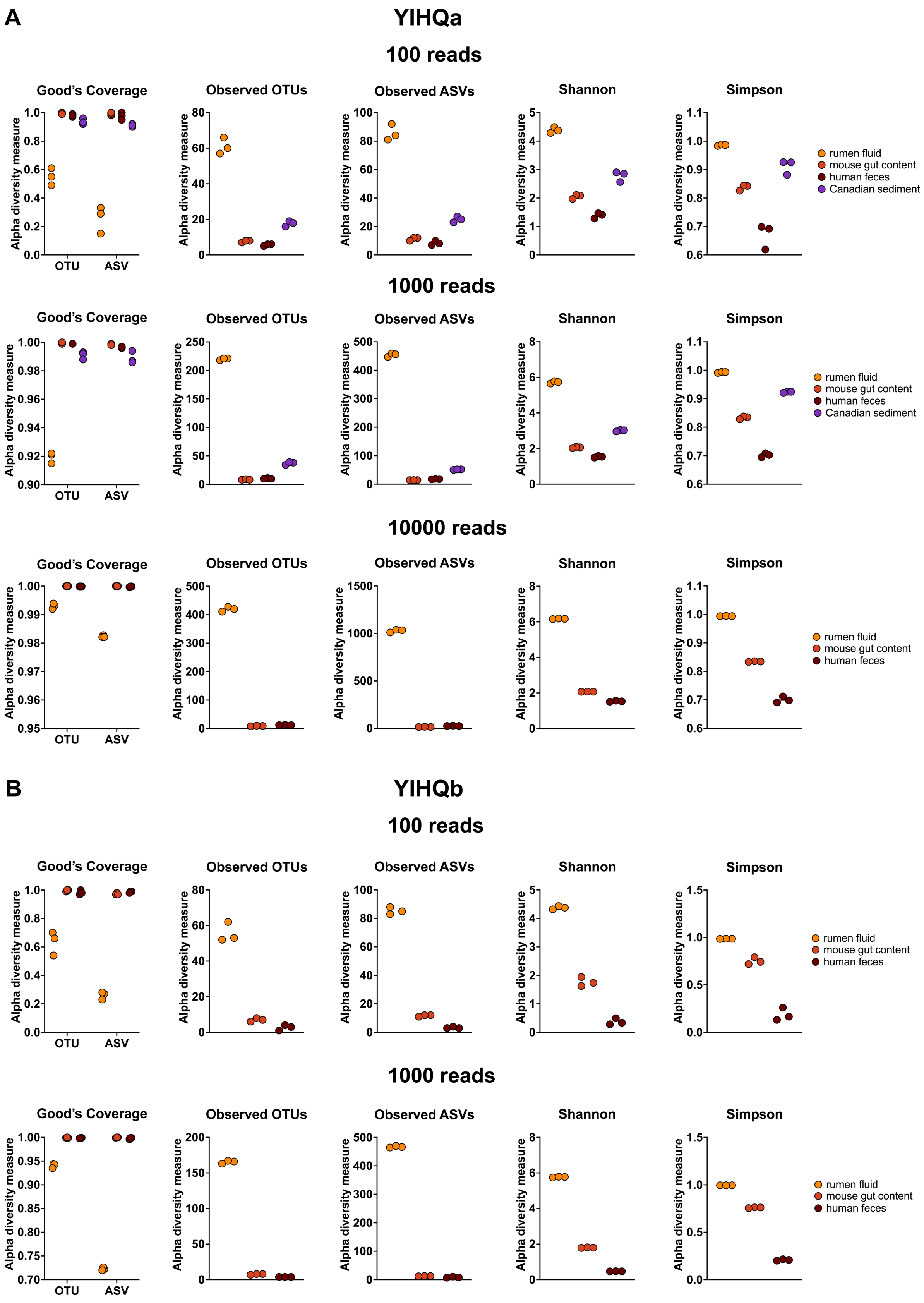
A
YIHQa
100 reads
Good's Coverage
Observed OTUs
Observed ASVs
Shannon
Simpson
Alpha diversity measure
OTU
ASV
rumen fluid
mouse gut content
human feces
Canadian sediment
1000 reads
10000 reads
B
YIHQb
100 reads
1000 reads

**Figure S3. Alpha diversity analysis of *yihQ* amplicon sequence data shows highly consistent results for all triplicates.**

Good's coverages, numbers of observed OTUs and ASVs, the Shannon diversity index, and the Simpson's diversity index were inferred from *yihQ* amplicon sequence data obtained with the YIHQa assay (**A**) and the YIHQb assay (**B**) Sequencing was performed in technical triplicates for each environmental and intestinal sample. The analysis was performed with verified *yihQ* sequences. Libraries were re-sampled at 100, 1000, and 10000 reads (Table S6).

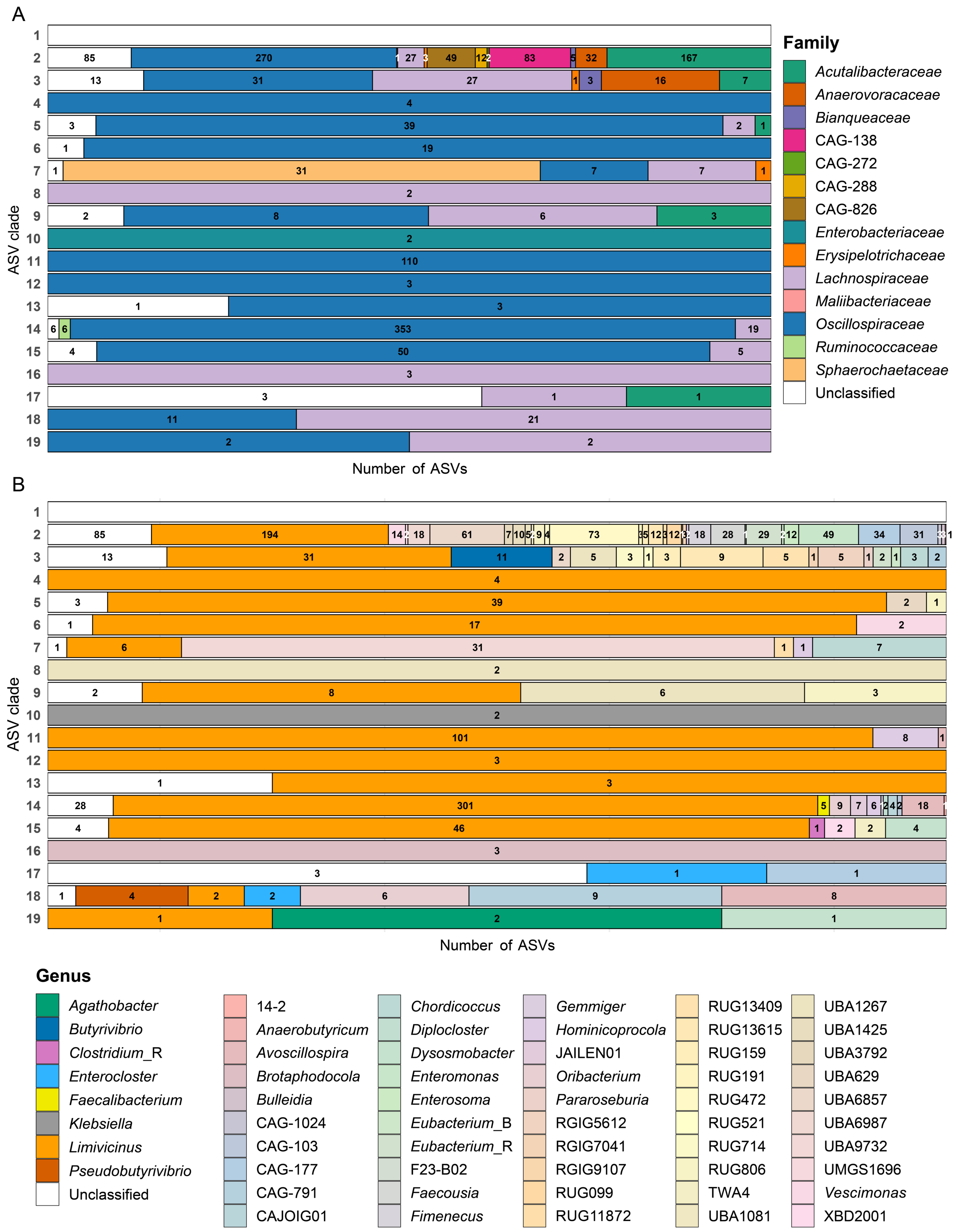

**Figure S4. Taxonomic assignment of environmental *yihQ* ASVs.**
Taxonomic composition of *yihQ* ASV clades from the YIHQb amplicon sequencing assay (Fig. 3) shown at the family (**A**) and genus level (**B**). Translated *yihQ* ASVs were queried against the GlobDB database using BlastP. The two top-scoring hits for each ASV were used for taxonomic assignment. In cases where two equally scoring hits had divergent taxonomic assignments, taxonomy was set to "unclassified" at the conflicting rank. ASVs with no match or matches

below 90% sequence identity were also categorized as unclassified (white). Each horizontal bar represents an ASV clade, as defined in Fig. 3A, with segments indicating the percentage of ASVs assigned to each taxon within the clade. Numbers in bar plots refer to the absolute number of ASVs per taxon within one ASV clade. Taxonomic assignment at the class level is presented in Fig. 3B. Full taxonomic details for individual ASVs are available in Table S7.

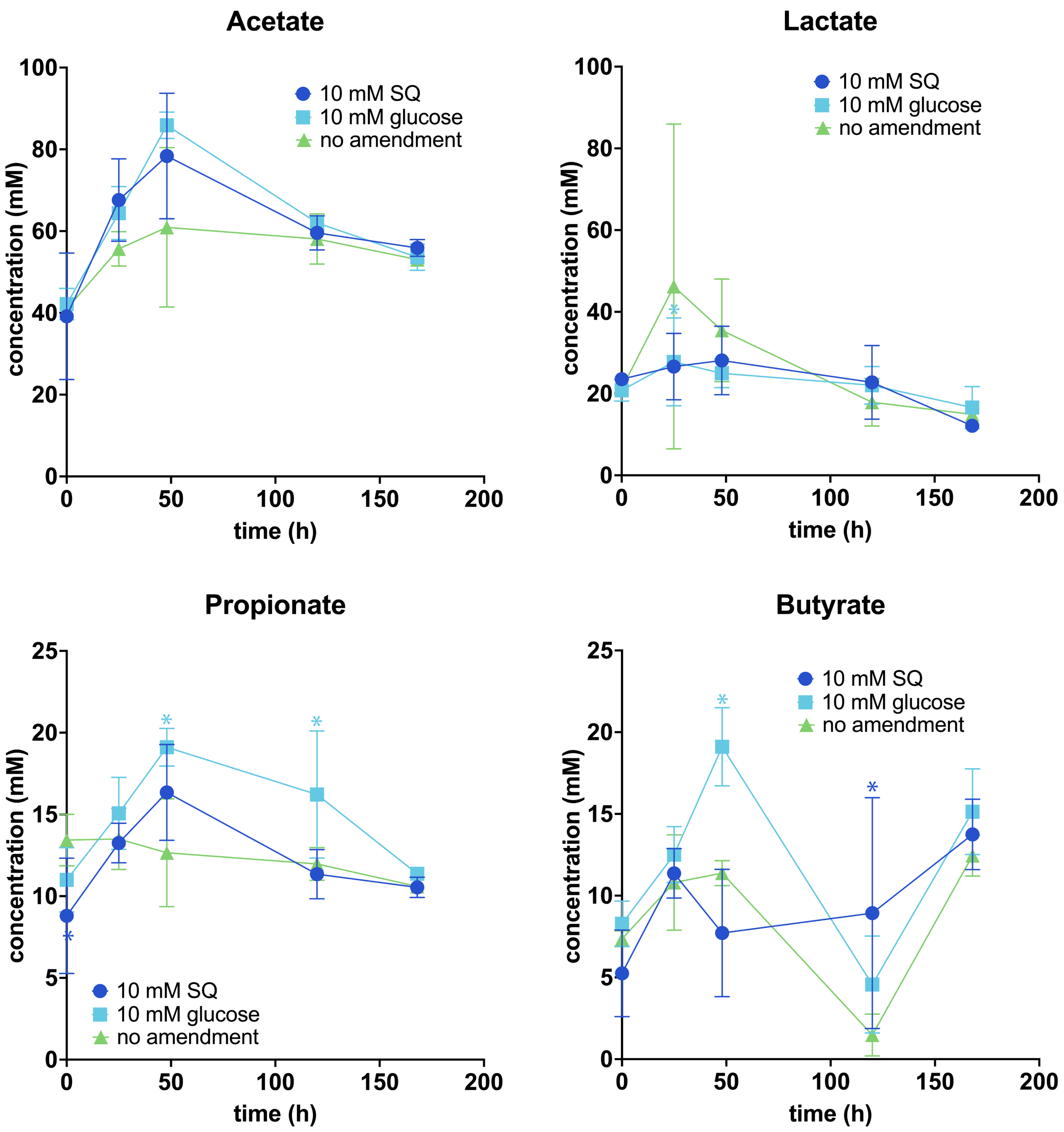

**Figure S5. Dynamics of volatile fatty acids in rumen fluid microcosms.**
Concentrations of volatile fatty acids are represented as the average of triplicate measurements. Error bars represent one standard deviation. Coloured asterisks show significant differences between the respective treatment and the unamended controls at individual time points. Significant differences were determined by ANOVA with Dunnett's post hoc test. Adjusted *p*-values of smaller than 0.05 were regarded as significant. SQ, sulfoquinovose.

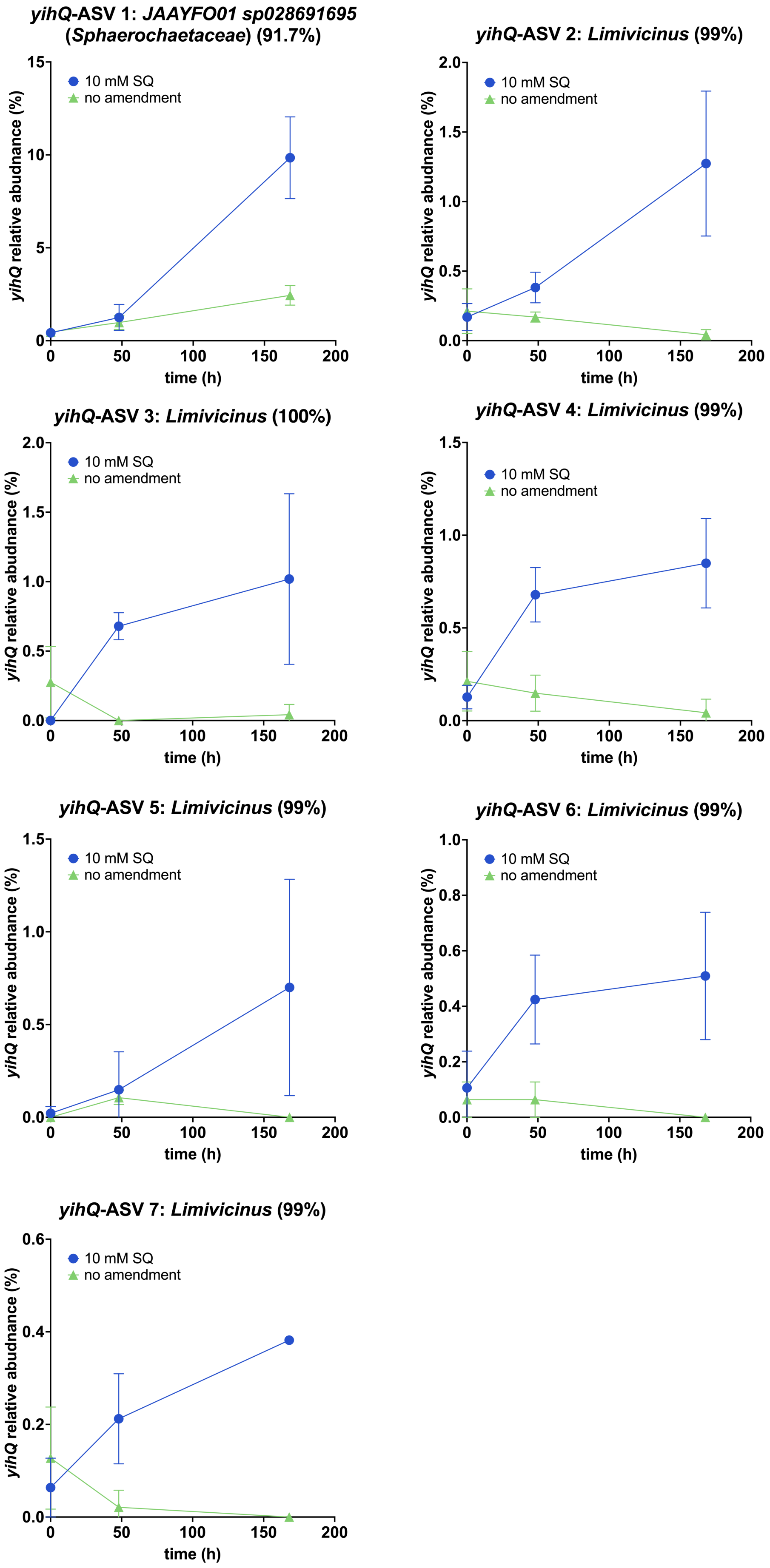
yihQ-ASV 1: JAAYFO01 sp028691695 (Sphaerochaetaceae) (91.7%)
yihQ-ASV 2: Limivicinus (99%)
yihQ-ASV 3: Limivicinus (100%)
yihQ-ASV 4: Limivicinus (99%)
yihQ-ASV 5: Limivicinus (99%)
yihQ-ASV 6: Limivicinus (99%)
yihQ-ASV 7: Limivicinus (99%)
10 mM SQ
no amendment
yihQ relative abudnance (%)
time (h)
0
50
100
150
200
0.0
0.2
0.4
0.5
0.6
0.8
1.0
1.5
2.0
5
10
15

**Figure S6. Relative abundance dynamics of *yihQ*-ASVs that were significantly enriched in SQ-amended rumen fluid microcosms.**

ASVs are shown in order of their average relative abundance increase compared to the unamended control. Points represent averages of three replicates and error bars represent one standard deviation. Taxonomic classification and the *yihQ* sequence similarity to the closest relative (in brackets) is shown for each ASV. ASV, amplicon sequencing variant; SQ, sulfoquinovose.
