## Additional files 1-7 for "Extensive richness and novel taxa of sulfoquinovose-degrading bacteria in the cow rumen": README.txt

Contents of Additional_file_1
• YIHQ_all_sequences.faa – Amino acid sequences of all retained homologs (n = 500).
• YIHQ_ingroup.id – Sequence IDs classified as the ingroup (n = 285).
• YIHQ_outgroup.id – Sequence IDs classified as the outgroup (n = 215).
• YIHQ_dereplicated_sequences.faa – Amino acid sequences of homologs after de-replication at 95% protein sequence identity (n = 294).
• YIHQ_dereplicated_AA_alignment.aln – MUSCLE alignment of de-replicated homologs (n = 294).
• yihQ_tree.nwk.nwk – YihQ reference tree inferred from de-replicated amino acid sequences (n = 294).
